# Forward variable selection improves the power of random forest for high-dimensional microbiome data

**DOI:** 10.1101/2020.10.29.361360

**Authors:** Tung Dang, Hirohisa Kishino

## Abstract

**Background:** Random forest (RF) captures complex feature patterns that differentiate groups of samples and is rapidly being adopted in microbiome studies. However, a major challenge is the high dimensionality of microbiome datasets. They include thousands of species or molecular functions of particular biological interest. This high dimensionality significantly reduces the power of random forest approaches for identifying true differences. The widely used Boruta algorithm iteratively removes features that are proved by a statistical test to be less relevant than random probes.

**Result:** We developed a massively parallel forward variable selection algorithm and coupled it with the RF classifier to maximize the predictive performance. The forward variable selection algorithm adds new variable to a set of selected variables as far as the prespecified criterion of predictive power is improved. At each step, the parameters of random forest are optimized. We demonstrated the performance of the proposed approach, which we named RF-FVS, by analyzing two published datasets from large-scale case-control studies: (i) 16S rRNA gene amplicon data for *Clostridioides difficile* infection (CDI) and (ii) shotgun metagenomics data for human colorectal cancer (CRC). The RF-FVS approach further screened the variables that the Boruta algorithm left and improved the accuracy of the random forest classifier from 81% to 99.01% for CDI and from 75.14% to 90.17% for CRC.

**Conclusion:** Valid variable selection is essential for the analysis of high-dimensional microbiota data. By adopting the Boruta algorithm for pre-screening of the variables, our proposed RF-FVS approach improves the accuracy of random forest significantly with minimum increase of computational burden. The procedure can be used to identify the functional profiles that differentiate samples between different conditions.

## Background

A microbiome is the full collection of genes of all microbes in a community; for example, all bacteria in a sample from the gut of a healthy individual or from an individual with a disease. Identifying deferent microbiome compositions between two or more groups is one of the most important purposes of microbiome studies [1, 2]. High-throughput sequencing technologies have allowed the microbiome composition and function in different environments to be quantified correctly [3, 4]. Several marker identification methods have been developed for applications in microbiome studies. The standard statistical approaches, such as Kruskal–Wallis (KW) test with the Benjamini–Hochberg false discovery rate (FDR) correction [5] or blocked (univariate) Wilcoxon tests [6], measure taxon relative abundances, analyze within- and between-sample diversity (α and β diversity, respectively), and perform classical hypothesis testing. These approaches are limited in their ability to classify unlabeled data or to extract salient features from highly complex and/or sparse datasets.

Machine learning technology has been applied in microbiome studies, especially for predicting specific diseases and supporting medical diagnosis [7, 8]. Because random forest (RF) captures the complex feature patterns that differentiate groups of samples [9, 10], it is rapidly being adopted for the analysis of microbiome data. The RF algorithm is a modification of bagging that aggregates a large collection of decision trees [11]. A main step in building an ensemble of decision trees is to perform random sampling of the available features to generate different subspaces of features at each node of each unpruned decision tree. This strategy can produce better estimation performances than a single decision tree because each tree estimator has low bias but high variance, whereas a bias-variance trade-off is achieved by the bagging process of RFs. RF methods have been applied successfully to genetic and microbiome data [12, 13, 14]. It is anticipated that RF methods and implemented importance measures will help in the identification of microbiome species that can be used to distinguish diseased and non-diseased samples. Identifying a core set of the most significant microbial species is of high interest, not only for diagnosis of certain diseases but also to gain valuable insights into the biological functionality and mechanisms of these species.

However, the performance and diversity of decision trees in the ensemble significantly influence the performance of RF algorithms. The generalization error for RFs involves measures of how accurate the individual classifiers are and their interdependence. Therefore, the high dimensionality problems of microbiome datasets pose a number of challenges. For example, microbiome datasets tend to contain a large number of microbiome species whose functions may not be related to the disease of interest. Common random sampling methods may select a sizeable number of subspaces that do not include the informative microbiome species and functions. As a consequence, the decision trees generated from these subspaces will have reduced average strength, thereby increasing the error bounds for the RF algorithm. A number of different approaches have been proposed to identify important variables that could improve the performance of RF algorithms. For example, the Boruta algorithm [15] was proposed to identify a set of relevant features using an RF classification algorithm that iteratively removes the variables using a statistical test. These relevant features are different from the objective of relevant and also non-redundant feature subsets. Moreover, a standard permutation test [16] was proposed to estimate the distribution of measured importance values of the RF algorithm for each predictor variable by repeatedly permuting the variable and randomly shuffling the data values so that the original association between the response and predictor variables was destroyed. A high value of the permutation importance of the predictor variable indicates high significant association to the response. However, the sizes of the selected subsets of features are still large in the high-dimensional microbiome database, the power of RF algorithm is not significantly improved, and it is difficult to interpret the selected features.

Although a number of different algorithms and tools have been developed for microbiome analysis [7], effective approaches require a lot of possible combinations of variables, which exponentially increases the computational burden as the number of involved features increases. Even though a small number of machine learning methods, including the RF algorithm, can be easily parallelized, building prediction models for thousands of microbiome species and functions can be very time consuming.

In this study, we propose a novel procedure that tackles the challenges described above. The core of our procedure is an RF classifier coupled with forward variable selection (RF-FVS), which selects a minimal-size core set of microbial species or functional signatures to maximize the predictive performance of the RF classifier. To reduce the computational cost, we designed a parallelized algorithm and integrated a prescreen algorithm. To examine the performance of the RF-FVS approach, we analyzed two empirical datasets from large-scale case control studies. One is a published case-control 16S rRNA gene amplicon sequencing gut microbiome dataset with 3347 operational taxonomic units (OTUs) for *Clostridioides difficile* infection (CDI) [17] from 338 individuals. The dataset included disease meta data and sequencing data for three groups: 89 individuals with CDI (cases), 89 with diarrhea who tested negative for CDI (diarrheal controls), and 155 non-diarrheal controls. The other is a fecal shotgun metagenomic dataset [18] that included 290 samples from tumor-free controls and 285 samples from individuals with colorectal cancer (CRC). The RF-FVS serves as the basis for the integrated analysis of microbiome data, detecting significant species in a phylogenetic tree [19] and predicting functional capabilities of microbial communities based on 16S rRNA datasets [20]. To estimate the dependence of the pipeline on the quality of the microbiome functional profile data, we analyzed the predicted functional profiles obtained from the 16S rRNA dataset, which were approximately 85% accurate, and the functional profiles obtained from the shotgun metagenomic dataset. We found that our RF-FVS performed well even for the predicted functional profile data.

## Results

### Improved accuracy for relative OTU abundance data from the 16S rRNA gene amplicon dataset

We compared the predictive power of our RF-FVS approach to classify three groups (CDI case, diarrheal control, non-diarrheal control) between the microbiome data and the clinical data. The clinical data included age, sex, ethnicity, antibiotic use, antacid use, a vegetarian diet, surgery within the past 6 months, a history of CDI, residence with another person who had CDI, and residence with another person who works in health care. The clinical data were 51% accurate (AUC = 0.7), whereas the microbiome data were 81% accurate (AUC = 0.95) (Figure S1). In particular, the accuracy was high for the microbiome data of the non-diarrheal control group. Moreover, when treating the composition of the microbiota data by Phylogenetic Isometric Log-Ratio (PhILR) transformation, we found that the RF classifier achieved high accuracy (85%; AUC = 0.95). However, it was still difficult to distinguish CDI cases and diarrheal controls in both the microbiome and phylogenetic transformation data. Overall, the accuracy of diagnosis was significantly improved with the RF-FVS approach for the CDI and diarrheal control groups for the microbiome data compared with the accuracy with the RF-PI and RF-BR algorithms (Fig. 1). By focusing on 119 species, the accuracy of the RF classifier increased to 96% (AUC = 0.99) (Figure S1). CDI cases were identified with an accuracy of 94% (AUC = 0.94) and diarrheal controls were identified with an accuracy of 93% (AUC = 0.97). For the phylogenetic transformation data, the accuracy was 95% (AUC = 0.97) when the FVS approach detected the 36 phylogenetic internal nodes.

**Fig 1.**
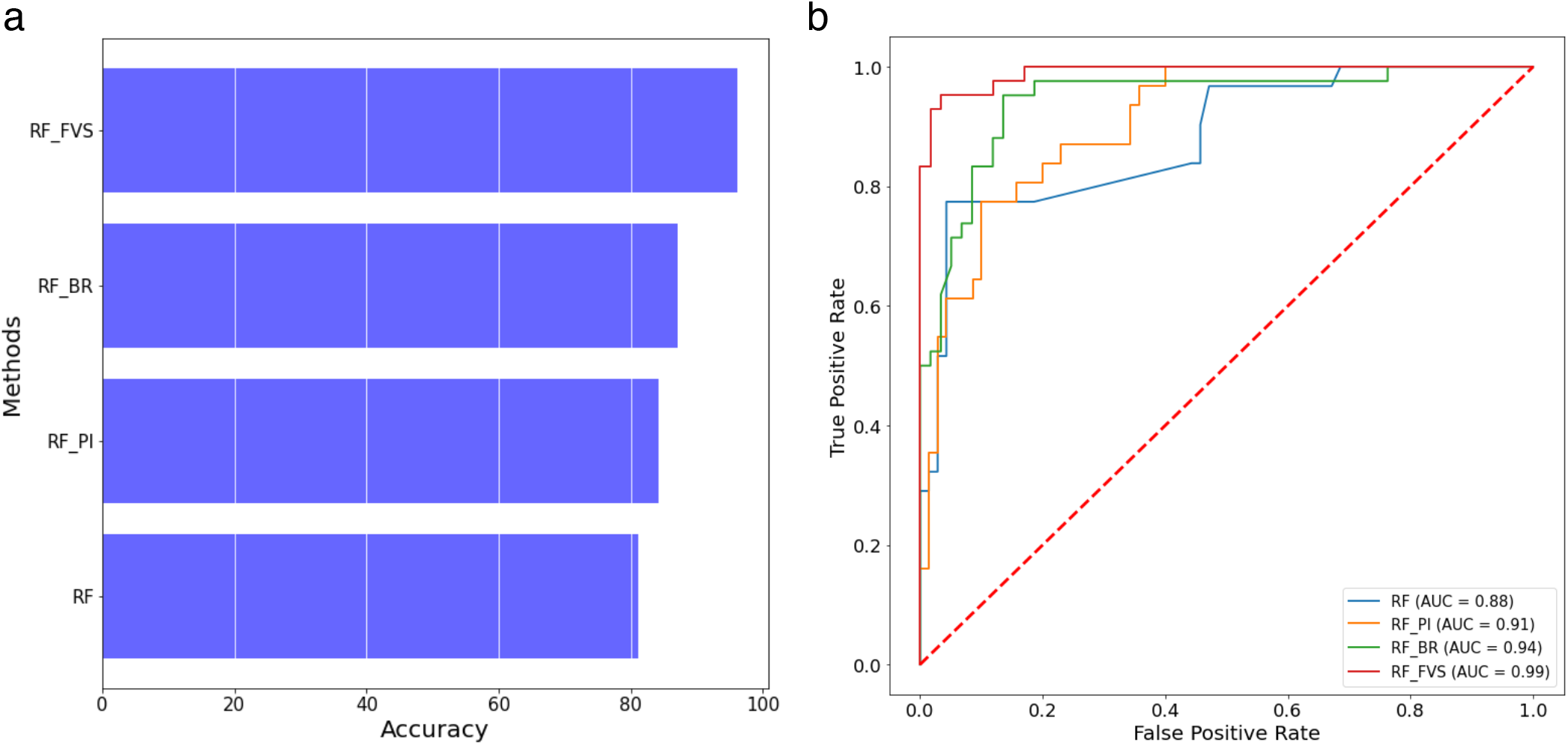
Performance of the random forest classifier for 16S rRNA gene amplicon data from the *Clostridioides difficile* infection (CDI) dataset. **a** Accuracy of random forest classifier with different variable selection methods for the 16S rRNA gene amplicon dataset. **B** ROC and AUC of random forest classifier for the 16S rRNA gene amplicon dataset; RF: Random forest algorithm, RF-PI: Random forest algorithm with permutation importance algorithm, RF-BR: Random forest algorithm with Boruta algorithm, RF-FVS: Random forest algorithm with forward variable selection algorithm.

### Mapping the selected species on the 16S rRNA phylogenetic tree

The forward variable selection approach avoided the sparse problems in selection of microbial species that the Boruta and permutation importance algorithms could not overcome (Fig. 2). Therefore, the number of selected species was significantly smaller than the numbers with the other methods, and the performance of the RF algorithm was improved and easier to interpret. The 119 selected OTUs from nine families were clustered in the 16S rRNA tree: *Peptostreptococcaceae, Verrucomicrobiaceae, Veillonellaceae, Ruminococcaceae, Lachnospiraceae, Bacteroides, Porphyromonadaceae, Lactobacillaceae*, and *Enterobacteriaceae* (Fig. 2a). They formed three main groups of species that were associated with the differentiation of CDI cases from the non-diarrheal controls. The species in group B in Fig. 2a (such as denovo 54, 1326, 601, 2346, and 3486) that showed strong positive correlations with CDI cases belonged to a diverse number of families (Figure S2 and Table S1) such as *Peptostreptococcaceae* [21], *Lactobacillaceae* (including *Lactobacillus* genus), *Enterococcaceae* (including *Enterococcus* genus) [22, 23], *Verrucomicrobiaceae*, and *Veillonellaceae* [24]. For example, species (such as denovo 54) in *Clostridium cluster XI* in family *Peptostreptococcaceae* that showed high positive correlations with CDI cases (Figure S2) are main candidates for treatment development [25] and the study of new challenges that have arisen because of the discontinuation of therapy for CDI disease [26]. Pérez-Cobas *et al*. [27] reported that the most striking changes in the microbiome of CDI cases occurred in the *Lactobacillaceae* family, whose frequency increased from <1% at the beginning of antibiotic treatment to 83.3% and 70% on days 35 and 38 of an antibiotic course, then reduced to 15.5% after antibiotic therapy. The species in group B that were selected by the Boruta and permutation importance algorithms, were similar to those selected using the forward variable selection approach (Fig. 2b and 2c). Most of the species in group C (such as denovo 127, 1399, and 788) that showed positive correlations with CDI cases belonged to the *Enterobacteriaceae* family (Fig. 2a and Table S1). Studies [28, 29, 30] have shown that relative overgrowth of members of the *Enterobacteriaceae* family was one of the main causes of significantly disturbed microbiota in CDI. Thus, *C. difficile* colonization may be facilitated by increased endotoxin production with increased intestinal permeability. However, in group C, the Boruta and permutation importance algorithms selected more species associated with CDI than the forward variable selection approach (Fig. 2b and 2c) because these two algorithms used the decrease of Gini impurity after a node split as the main input data for computational processes in order to select the main features. The corresponding species that became the potential candidates of these two algorithms showed large decreases of impurity after certain split. The reduction of impurity of species became very slow and the differences of impurity between species in the same families were insignificant in the high-dimensional sparse microbial data (Figure S3 and Table S1). Therefore, the abilities of the Boruta and permutation importance algorithms were influenced significantly even if the statistical tests were used. For example, in group C, the FVS algorithm selected a few species in *Lachnospiraceae* family that had positively associated OTUs (Fig. 2a), but the other algorithms kept a large number of the species in this family that showed poor positive correlations with CDI cases (Fig. 2b, 2c and Table S2). In groups B and C, a large number of species in the *Lachnospiraceae* family that were kept by the Boruta algorithm, were associated with non-diarrheal controls (Fig. 2b) but showed poor correlations with non-diarrheal controls (Table S2). Besides, a number of species in the *Ruminococcaceae* family that were selected only by the Boruta and permutation importance algorithms, showed poor positive correlations with non-diarrheal controls (Fig. 2b, 2c and Table S3).

**Fig 2.**
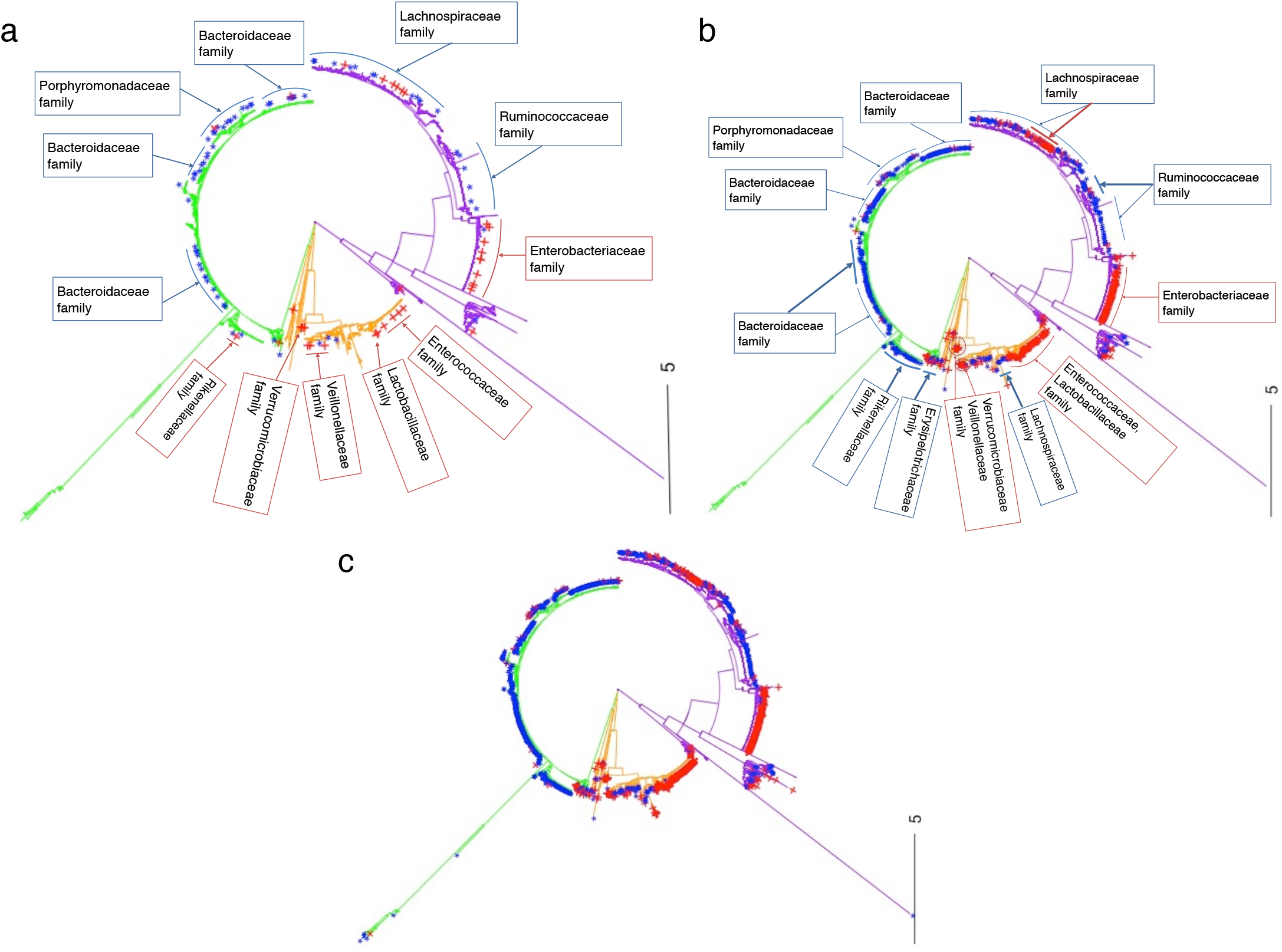
Phylogenetic tree of the 16S rRNA microbiota (OTU) data from the CDI dataset. **a** Positions indicate the 119 important microbial species that remained after forward variable selection. ***b*** Positions indicate about 1000 important microbial species that remained after applying the Boruta algorithm. **c** Positions indicate about 2000 important microbial species that remained after applying the permutation importance algorithm. Red indicates OTUs associated with CDI cases; blue indicates OTUs associated with the non-diarrheal controls; green indicates group A; orange indicates group B; purple indicates group C.

Most species in group A (such as denovo 557, 302, 1888, 1987, 1983, and 610), some species in group C (such as denovo 26, 9, 1295, 447, and 347) and one species in group B (denovo 156) that were enriched in the non-diarrheal controls belonged to the *Ruminococcaceae, Lachnospiraceae, Bacteroides*, and *Porphyromonadaceae* families (Fig. 2a, Figure S2 and Table S1). Short-chain fatty acid (SCFA) production is known to play a principal role in the regulation of intestinal inflammatory processes [31] and intestinal barrier maintenance [32]. CDI was found to cause significant reductions in the *Ruminococcaceae* and/or *Lachnospiraceae* families that produce butyrate and SCFA [33, 34]. *Lachnospiraceae* and *Ruminococcaceae* sequences were dominant at 45.8% and 17.4% respectively in healthy fecal microbiota, whereas sequences from other families constituted <3% of all the pyrosequencing reads [34]. However, these proportions decreased significantly in the CDI group (*Lachnospiraceae* (11.2%), *Ruminococcaceae* (3.0%)) [34]. Moreover, the four species in group A (denovo 2019, 1773, 3205, and 2528) (Fig. 2a and Table S1) showed weak associations with CDI case. However, a significantly larger number of species, that had positive correlations with CDI, were selected by the Boruta and permutation importance algorithms (Fig. 2b and 2c). For example, in group A, the number species in the *Bacteroidaceae* family that were kept only by these algorithms, showed insignificant positive correlations with non-diarrheal controls (Table S4). Moreover, the Boruta and permutation importance algorithms selected a huge number of species in the *Rikenellaceae* and *Erysipelotrichaceae* families that were ignored by the FVS approach (Fig. 2); however, they had very poor correlations with CDI case and non-diarrheal controls (Table S5 and S6).

When we applied our RF classifier to the clinical data, we found that antibiotic treatment contributed significantly to an increase in the accuracy of the prediction models [27, 35].

### Improved accuracy for relative OTU abundance data from shotgun metagenomics data

Our RF classifier successfully distinguished the CRC cases and tumor-free controls in both the 16S rRNA gene amplicon data and the shotgun metagenomics data (Figure S4). The average accuracy for the 16S rRNA gene amplicon data, which included 18,448 OTUs, was 68.18% (AUC = 0.73). For the shotgun metagenomics data, the accuracies of the RF classifier for CRC cases and tumor-free controls were 70% (AUC = 0.86) and 80% (AUC = 0.86) respectively. The forward variable selection significantly improved the performance of the RF algorithm as shown in Fig. 3. Specifically, the forward variable selection detected 75 microbial species (out of 849 species) that were differentially abundant in the CRC microbiome, which increased the accuracy of the RF classifier to 88% (AUC = 0.92) for the CRC cases and to 95% (AUC = 0.92) for the tumor-free controls (Figure S4). These findings are consistent with previous reports of significant enrichment of novel species in the fecal microbiomes of patients with CRC. For example, we detected the three most important species, *Parvimonas micra, Flavonifractor plautii*, and *Gemella morbillorum*, that helped to improve the accuracy of our RF classifier (Figure S5). Yu *et al*. [36] and Feng *et al*. [37] identified *Parvimonas micra* as a key species that was consistently enriched in the microbiomes of patients with CRC, and we found this species had the most importance among our selected species. Gupta *et al*. [38] found that *Flavonifractor plautii* was associated significantly and enriched in CRC samples of Indian patients. *Flavonifractor plautii* was linked with the degradation of beneficial anticarcinogenic flavonoids, and this role was strongly correlated with enzymes and modules involved in flavonoid degradation in CRC samples of the Indian patents. In our study, *Flavonifractor plautii* was significantly associated with CRC samples of cohorts from France, Germany, China, United States, and Austria (Figure S6). Our analysis showed that the dysbiosis of fecal microbiota was characterized by enrichment of potential pathogens and the reduction in butyrate-producing members such as *Fusobacterium* in CRC samples [39, 40]. We also identified other main species (Figure S5), *Gemella morbillorum* [41], *Prevotella intermedia* [42], and *Prevotella nigrescens* [43], which have been shown previously to be strongly associated with an increased risk of CRC. In the gut microbiome of the tumor-free controls, we detected *Bifidobacterium adolescentis* and, *Lactobacillus ruminis*, which previously have been associated with healthy samples and negatively correlated with CRC cases (Figure S5). These species may have a protective role against the development of non-alcoholic fatty liver disease and obesity [44].

**Fig 3.**
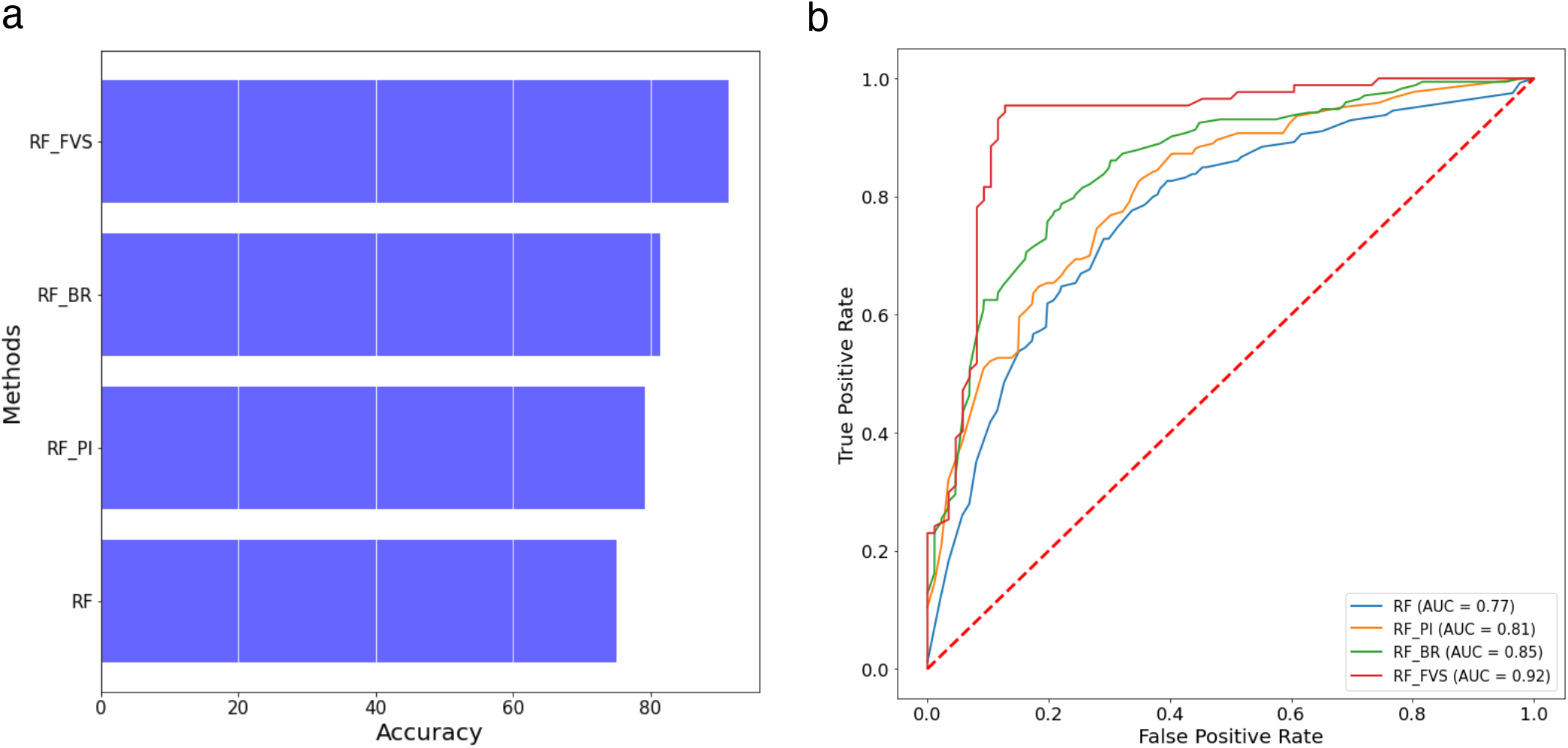
Performance of the random forest classifier for the shotgun metagenomics data of colorectal cancer (CRC) dataset. **a** Accuracy of random forest classifier with the different variable selection methods for the shotgun metagenomics dataset; **b** ROC and AUC of random forest classifier for the shotgun metagenomics dataset. RF: Random forest algorithm, RF-PI: Random forest algorithm with permutation importance algorithm, RF-BR: Random forest algorithm with Boruta algorithm, RF-FVS: Random forest algorithm with forward variable selection algorithm.

### Influential bacterial functions predicted from the 16S rRNA microbiota (OUT) data

The forward variable selection algorithm detected 119 OTUs out of 3347 OTUs. To predict functional profiles from these OTU candidates, we examined 5818 of the 21,620 functional profiles of the KEGG organisms in the Tax4Fun framework.

The RF method gave an average accuracy of 81% (AUC = 0.93) for the 5818 predicted functional profiles. Specifically, the accuracies for the CDI cases, diarrheal controls, and non-diarrheal controls were 68%, 79%, and 93% (AUCs = 0.88, 0.92, and 0.97) respectively. To reduce the computational burden, we used the prescreen algorithm at the first step, which reduced the predicted functional profiles from 5818 to 2534. Then, we applied the RF-FVS algorithm, which identified 23 functional profiles (out of 2534) that were different for the CDI cases compared with the controls, which significantly increased the average accuracy to 90% (AUC = 0.95) (Figure S7). Specifically, the accuracies for CDI cases, diarrheal controls, and non-diarrheal controls were 77%, 93%, and 98% (AUCs = 0.90, 0.95, and 0.97) respectively. Some of the 23 most significant functional profiles were strongly associated with CDI cases but a larger number of them were associated with healthy gut (non-diarrheal controls) (Fig. 4). Our results confirmed that bacteria support human health with functions such as histidinol dehydrogenase, gluconate 5-dehydrogenase, lactaldehyde reductase, and alpha-1,3-rhamnosyltransferase [45] [46, 47]. However, these functions were absent in the microbiomes of patients with CDI, mainly because they were treated with antibiotics and proton pump inhibitors, which killed these bacteria (Fig. 4). Therefore, *C. difficile*, which is resistant to these treatments, became dominant and increased the risk of CDI. Conversely, some of the functional profiles were strongly enriched in the CDI cases. They included alcohol dehydrogenase, L-iditol 2-dehydrogenase, and glucose 6-dehydrogenase, which are related to the growth of *C. difficile* [48, 49], and xanthine dehydrogenase, which is related to the high-level resistance of *C. difficile* to antibiotics treatments [50].

**Fig 4.**
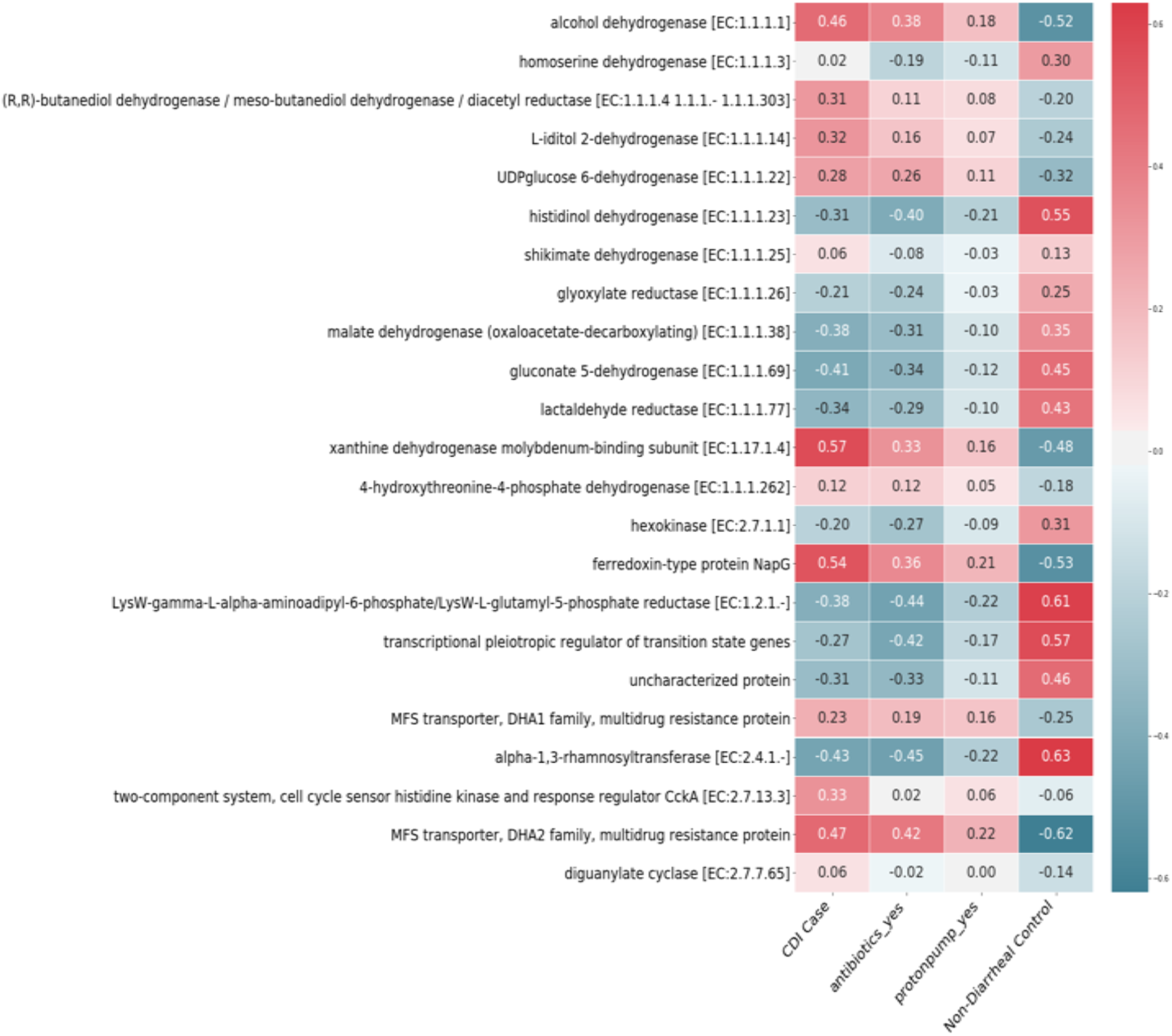
Correlations among the 23 main functional profiles with CDI cases, antibiotics, proton pump inhibitors, and non-diarrheal controls.

### Influential bacterial functions predicted using the shotgun metagenomics data

We used the evolutionary genealogy of genes from the Non-supervised Orthologous Groups (eggNOG) orthologous gene family abundances and KEGG module abundance profiles to detect functional profiles in the shotgun metagenomics data for CRC. Because the numbers of functions in the KEGG and eggNOG databases were very large (7955 and 31,185 respectively), we used the prescreen algorithm before applying forward variable selection.

Our RF-FVS algorithm identified 29 out of 7955 functions in the KEGG database that significantly improved the performance of the RF classifier. Specifically, the accuracy increased from 60% (AUC = 0.79) to 84% (AUC = 0.87) for CRC cases and from 80% (AUC = 0.79) to 97% (AUC = 0.87) for tumor-free controls (Figure S8). A number of functions such as HOMODA hydrolase (K10623), carbamoyl-phosphate synthase 1 (K01948), heptose-I-phosphate ethanolaminephosphotransferase (K19353), tartronate-semialdehyde synthase (K01608), ABC transporters (K11707), and biofilm formation - *Pseudomonas aeruginosa*/bacterial secretion system (K11903) had strong positive correlations with CRC cases (Figure S9). The contributions of some of these functions to the stage progress of CRC have been reported. For example, carbamoyl-phosphate synthase 1, a metabolic enzyme that utilizes ammonia to produce carbamoyl phosphate, is encoded by one of four novel driver genes that were identified as hubs for stage-III progression of colorectal cancer [51].

Our RF-FVS algorithm also identified 53 out of 31,185 functions in the eggNOG database that significantly improved the performance of the RF classifier. Specifically, the accuracy increased from 62% (AUC = 0.78) to 86% (AUC = 0.90) for CRC cases and from 80% (AUC = 0.78) to 97% (AUC = 0.90) for tumor-free controls (Figure S10). Although a large number of functions were significantly positively correlated with CRC cases, such as ENOG410Y6BY, ENOG410XYS8, ENOG411EMB, and ENOG410ZGTS (Figure S11), experimental information about their functions is lacking. These genes are likely to be good candidates for further studies.

### Forward variable selection and the prescreening algorithm reduced the CPU time

Although the RF classifier achieved high accuracy in analyzing the microbiome data, the high dimensionality of the data meant the computational burden was high. The RF classifier took about one week to identify 119 species out of 3347 species in the 16S rRNA gene amplicon data for CDI. By using Boruta algorithm to identify a small number of informative variables, the 3347 species were reduced to 1008 species and the RF-FVS algorithm detected 96 species with an increased accuracy of 99% (AUC = 0.99) (Table 1). The computation time also was reduced from one week to one day. For the functional profile predictions, the FS-FVS algorithm took about 4 days to identify 65 out of 5818 functional profiles. The Boruta algorithm reduced the total number of functional profiles from 5818 to 2534 and the RF-FVS then needed only 13 hours to detect 23 out of the 2534 functional profiles. The accuracy of the RF classifier increased to 90.1%.

**Table 1.** Random forest classifier with forward variable selection (RF-FVS) with and without the prescreen algorithm for the CDI dataset. ✓, the prescreen algorithm was used; ✗, the prescreen algorithm was not used. All algorithms were run in a parallel environment. The properties of the parallel version were evaluated on a high-performance computer (Intel^®^ Xeon^®^ Gold 6230 Processor 2.10 GHz × 2, 40 cores, 2 threads per core, 93.1-Gb RAM) under Ubuntu 20.04.1 LTS.

| Data type | Analysis framework |  |  | Accuracy | Time computation |
| --- | --- | --- | --- | --- | --- |
|  | Boruta algorithm | Random forest | Forward algorithm |  |  |
| OTUs data | ✗ | ✓ | ✓ | 96.04% | 1 week |
|  | ✓ | ✓ | ✓ | 99.01% | 1 day |
| PhILR transformation | ✗ | ✓ | ✓ | 95.05% | 26 hours |
|  | ✓ | ✓ | ✓ | 95.05% | 8.75 hours |
| Functional profile data | ✗ | ✓ | ✓ | 90.1% | 4 days |
|  | ✓ | ✓ | ✓ | 90.1% | 13 hours |

## Discussion

A number of publicly available databases contain information about microbial species and their functions that are associated with disease or health. The large number of microbial species and functional profiles in these databases significantly negatively influence the power of machine learning classifiers, including the RF algorithm. Further, the computational cost of running these algorithms to detect a few informative features in the high-dimensional space of microbiome data is still very high. In this study, we developed a novel procedure that significantly enhances the RF classifier and substantially improves its performance in terms of high-speed computation and high accuracy. We tested its performance using two microbiome datasets that contained a large number of species and functional signatures (>30,000 variables) but a very small proportion of significant variables.

Our RF-FVS approach was useful in several respects. Firstly, the RF-FVS algorithm identified the core set of microbial species and their functional profiles, which considerably increased the predictive accuracy of the RF classifier. The highest increase in the predictive accuracy was to 99% for the CDI cases classification. Moreover, because 16S rRNA sequencing data do not directly provide insights into the functional capabilities of the microbiome community, we integrated the Tax4Fun tool into our pipeline to predict the functional profiles of microbial communities based on the 16S rRNA datasets. Therefore, our RF-FVS approach could detect the minimal-size core set and optimal predictive subset of functional profiles of the selected microbial species and the linkages among them, which will provide insights into the ecological functioning of habitats. Some unknown species and genes within the dedicated taxa and functional profiles were detected in the group of selected features that made meaningful contributions to the predictive performance of the RF classifier. These species and genes are likely to be good candidates for future experimental studies.

Secondly, there are a number of standard approaches that can use 16S rRNA data to infer the metabolic potential of the corresponding microbial species. For example, if the database is annotated by the Greengenes database, PICRUSt can achieve good estimations for the functional potential of microbial communities [52]. Tax4Fun is a good option for data annotated by the SILVA database. In this study, the published databases that we used to checked the performance of the RF-FVS approach adopted the Ribosomal Database Project (RDP) approach to classify the 16S rRNA gene sequences taxonomically [53]. Tax4Fun achieved significantly better quality of predicted functional profiles than PICRUSt; therefore, Tax4Fun was integrated as the main option in our pipeline. In the future, we plan to check the performance of the RF-FVS approach for other databases that provide taxonomy annotations, such as Greengenes, LTP, RDP, and SILVA [54].

Moreover, to overcome the computational burden of high-dimensional data that limits the implementation of existing machine learning approaches, we developed a parallel computational strategy algorithm for handling large-scale problems in the forward variable selection algorithm. This parallel strategy helps to equally divide the computational burden of search processes among processors. Thus, the forward variable selection computation process is completely optimized and parallelized based on data partitioning. In microbiome datasets of tens of thousands of species and their functional profiles, selecting only a few hundred of the most significant samples can be a major problem. Our current strategy is focused on parallelly searching for the features of interest. In the future, the numbers of samples and features in microbiome datasets are likely to explode at a rapid pace. We anticipate that hybrid-partitioning strategies that partition the data both horizontally (over samples) and vertically (over features) will become essential to speed up the computational processes [55, 56].

## Methods

### Forward variable selection for random forests

The RF approach is an ensemble method that combines a large number of individual binary decision trees. Two main randomization procedures have been implemented to reduce variance of individual decision trees, deliver diversity amongst decision trees, and thus improve prediction accuracy. First, randomly selected training samples for each of the individual trees are applied to construct sufficiently diverse trees. Second, at each node within a tree, a set of randomly chosen candidate predictor variables is identified for the split.

However, random feature subspace sampling may not be a good strategy to deal with high-dimensional data because a large proportion of the features may not be informative of the class of an object in the high-dimensional data. If a random sampling strategy is implemented to select the subset of eligible features at each node, almost all the subsets are likely to contain a large number of non-informative features. For example, the 16S rRNA gene amplicon data for CDI [17] that we used to evaluate the performance of our proposed approach contained a total of 3347 microbiome species, but only 96 of the species were informative. Therefore, if a subset of species, which is usually the square root of the total number of species, is selected by resampling randomly at any node within the decision tree, the mean number of informative species selected at each node will be two. Therefore, individual decision trees built using such nodes will have low accuracy and the performance of the RF algorithm will suffer. In our approach, we used forward variable selection to identify a small number of informative variables to improve the performance of individual decision trees in an ensemble.

A key idea behind our algorithm was to divide the total number of variables into two groups, a remaining variables group and a selected variables group. We started with an empty group for the selected variables. At each step, a variable from the remaining variables group was added to the selected variables group such that the specified criterion was improved (i.e., area under the receiver operating characteristic [ROC] curve [AUC], a weighted average of the precision and recall [F1 score] or predictive accuracy). Model selection for microbial signature identification also can be performed using our RF-FVS algorithm. At each forward iteration, given the selected variables, the randomized parameter optimization algorithm for RF implements a randomized search over parameters, where each setting is sampled from a distribution over possible parameter values. Thus, the best RF model is specified by these selected variables.

Moreover, a high-speed computational strategy based on multi-processing architecture was developed to parallelize the forward variable selection algorithm at the single machine level and thus significantly reduce runtimes. Another key idea behind our algorithm was to create many subsets of variables, so that each subset had one of the variables from the remaining variables group added to the selected variables group. Because of the high dimensionality problems of microbiome data, the number of these subsets is usually significantly larger than the number of processors in a single computer system. Our solution was to create queues so that subsets are assigned randomly and each processor runs the computational processes from its own privately prepared queue. Fig. 5 shows how all computational burdens for searching important feature relevance are appropriately decomposed, so that they can be computed in a parallel environment. The RF model with the highest accuracy value for each subset of features is selected by a specific processor. Processors in symmetric multiprocessing communicate with each other through shared memory architecture to decide only the best feature among multiple candidates that should be added to the selected features. The algorithm stops when there are no additional variables that improve the current optimization parameters or when the maximum number of components to be included in the group of selected variables is achieved. The main parameters of the RF classifier that were optimized in our algorithm, were number of trees in the forest, maximum depth of the tree, minimum number of samples required to split an internal node, minimum number of samples required to be at a leaf node, and number of features to consider when looking for the best split.

**Fig 5.**
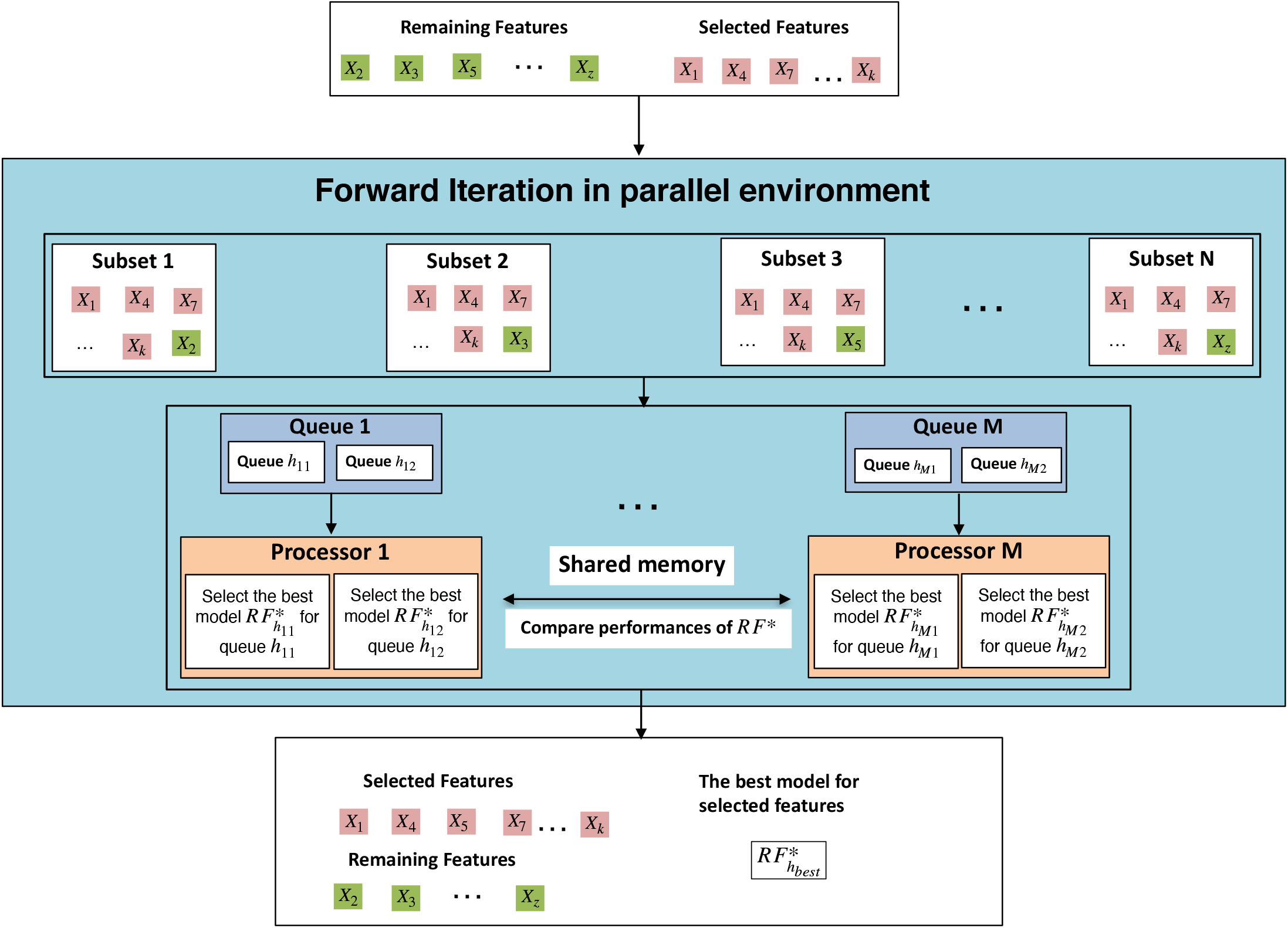
Massively parallel forward variable selection algorithm for the random forest (RF) classifier. The number of queues depends on the number of CPU cores available in the computer system.

### Functional gene enrichment analysis

Functional profiles were predicted from the 16S rRNA gene data for CDI using Tax4Fun. which was developed to analyze the enrichment of functional genes of microbiomes [20, 57]. The output from the QIIME software application with a SILVA database extension (SILVA 119) [58] was used to pre-process raw data for Tax4Fun. Tax4Fun transforms the SILVA-based operational taxonomic units (OTUs) into a taxonomic profile of KEGG organisms that is normalized by the 16S rRNA copy number (obtained from the NCBI genome annotations) [59]. The result is a table containing relative KEGG ortholog (KO) abundance levels. The KO profiles obtained from predictions or actually measured from shotgun metagenomic or metatranscriptomic data can be used for functional diversity profiling based on the KEGG annotation systems (pathways, modules, or EC categories). Because one KO can be assigned to multiple groups, our approach detects the most important functional profiles associated with disease or healthy samples.

### Phylogenetic transformation of microbiota data for random forest

To avoid spurious statistical analyses because of the relative nature of microbial abundance data in microbiota studies [60], we used the Phylogenetic Isometric Log-Ratio (PhILR) transformation [19]. The main idea behind the PhILR transformation is to consider the bacterial phylogenetic tree as a natural and informative sequential binary partition to construct an isometric log-ratio that converts compositional data into a real Euclidean space. This phylogenetically driven isometric log-ratio transformation can help to capture the hierarchical pattern of a microbial community structure. Therefore, the RF algorithm can identify the main internal nodes of a phylogenetic tree that represent phylogenetically-related bacterial groups (clades), thereby offering the opportunities for biological insights.

### Pre-screening algorithm for random forest coupled with forward variable selection

We used the Boruta algorithm [15] as a relevant embedded feature selection algorithm that uses the RF classifier to detect all strongly and weakly relevant OTUs (or phylogenetic internal nodes, or functional profiles) to reduce the considerable data dimensionality. This improved the classification accuracies and significantly decreased the time computation. The main idea of this algorithm was to duplicate each OTU, thus creating “shadow” OTUs by randomly permuting the observations of duplicated OTUs at the first step. Then, the importance of all the OTUs is computed (calculated as Z-scores) and the maximum Z-score among the shadow bands is identified when the RF classifier is run. The number of times that the importance of an OTU is higher than the maximum Z-score among the shadow OTUs is counted. An OTU is deemed “important” when the frequency is significantly higher than the expected value, otherwise the OTU is deemed “unimportant” and removed. Basically, the Boruta algorithm works with Scikit-learn dependency and acts as an interface medium on the RF classifier, which uses the importance measure generated by the original algorithm. The Boruta algorithm depends on the implementation of the Scikit-learn RF classifier, which runs on a single code to identify all the important features in a dataset with respect to an outcome variable. The global framework of our new approach is shown in Figure S12.

### Two empirical datasets

To examine the performance of our RF-FVS approach, we analyzed two empirical datasets from large-scale case control studies as follows.

#### *Clostridioides difficile* infection

We collected a published case-control 16S rRNA gene amplicon sequencing gut microbiome dataset that included disease meta data and sequencing data with 3347 OTUs for *Clostridioides difficile* infection (CDI) [17] from 338 individuals; 89 with CDI (cases), 89 with diarrhea who tested negative for CDI (diarrheal controls), and 155 non-diarrheal controls. To understand how clinical- and microbiome-based factors are associated with CDI, the gut microbiomes of these individuals were characterized. A total of 183 diarrheal stool samples from the 94 individuals with CDI, 89 diarrheal control samples, and 155 non-diarrheal control stool samples were analyzed. Taxonomic assignments were generated using a naive Bayesian classifier trained against a 16S rRNA gene training set provided by the Ribosomal Database Project (RDP) [53]. We also collected a set of published 16S rRNA gene sequences [61], among which, 120 samples were from individuals with human colorectal cancer (CRC) and 172 samples were from tumor-free controls. Taxonomic assignments were generated using the Bayesian classifier as described above.

#### Human colorectal cancer

We collected a published fecal shotgun metagenomic dataset [18] that included 290 samples from tumor-free controls and 285 samples from individuals with CRC. Taxonomic profiles were generated using the mOTU profiler tool [62] to guarantee consistency of the biological information. Unlike the 16S rRNA gene amplicon data, one of the most important advantages of fecal shotgun metagenomic data is that it provides functional abundance profiles, such as those for the eggNOG gene family or KEGG module abundance profiles, that allowed a direct analysis of the functional potential of the gut microbiome to be conducted.

## Availability of data and materials

RF-FVS is implemented in Python and is available on github (https://github.com/tungtokyo1108/Random_Forests_for_High-Dimensional_Data)

## Acknowledgment

We thank Margaret Biswas, PhD, from Edanz Group (https://en-author-services.edanz.com/ac) for editing a draft of this manuscript.

