## Supplemental information for "Forward variable selection improves the power of random forest for high-dimensional microbiome data"

### Supplementary Information

#### Supplementary Figure legends

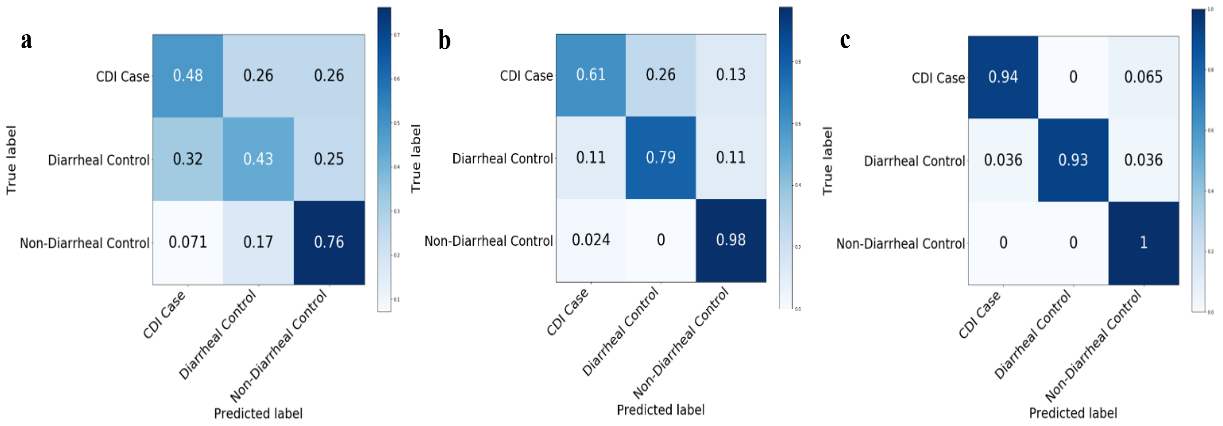


#### Additional file 1: Figure S1

Confusion matrices comparing the performance of the random forest classifier (with and without forward variable selection) for clinical data and 16S rRNA gene amplicon data of *Clostridioides difficile* infection (CDI) dataset. **a** Random forest classifier for the clinical data; **b** Random forest classifier for the 16S rRNA gene amplicon dataset; **c** Random forest classifier coupled with forward variable selection (RF-FVS) approach for the 16S rRNA gene amplicon dataset.

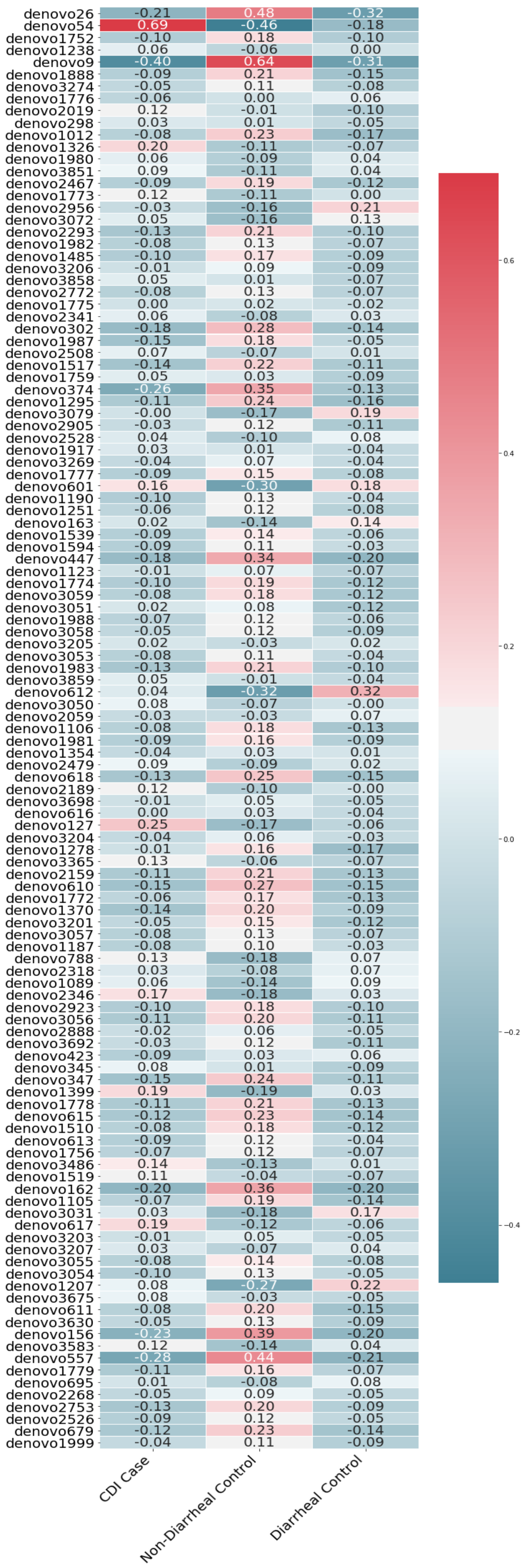


**Additional file 2: Figure S2a** Correlation among the 119 selected species with CDI cases, diarrheal controls, and non-diarrheal controls

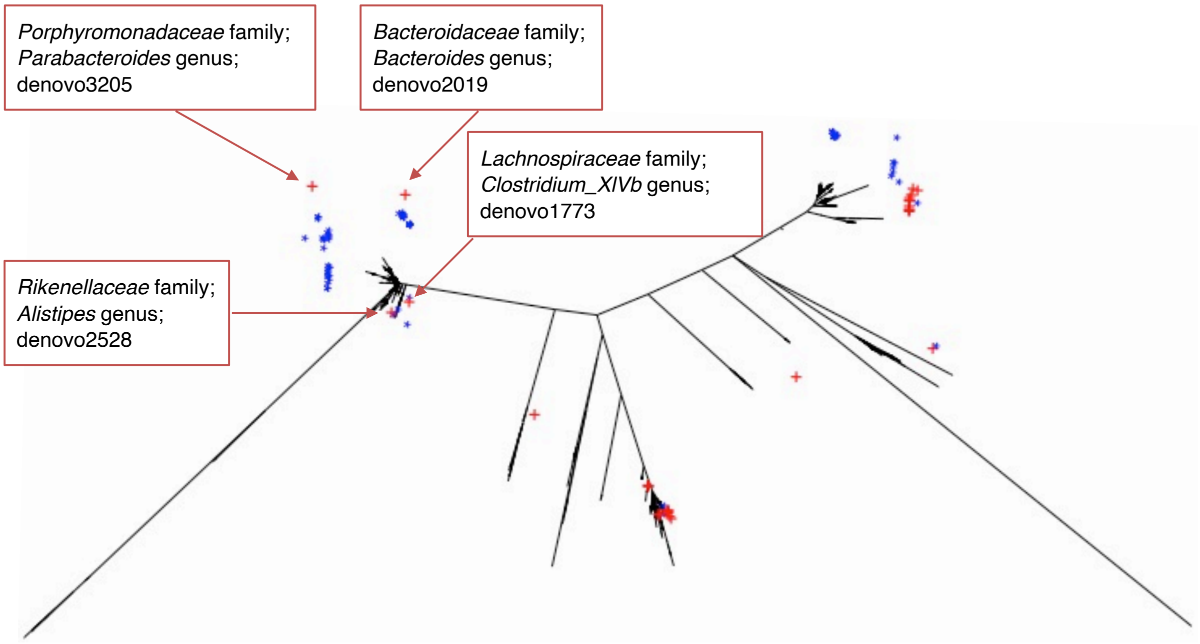


**Additional file 2: Figure S2 b**

Selected species in group A mapped on the 16S rRNA phylogenetic tree. Family and genus level information is shown.

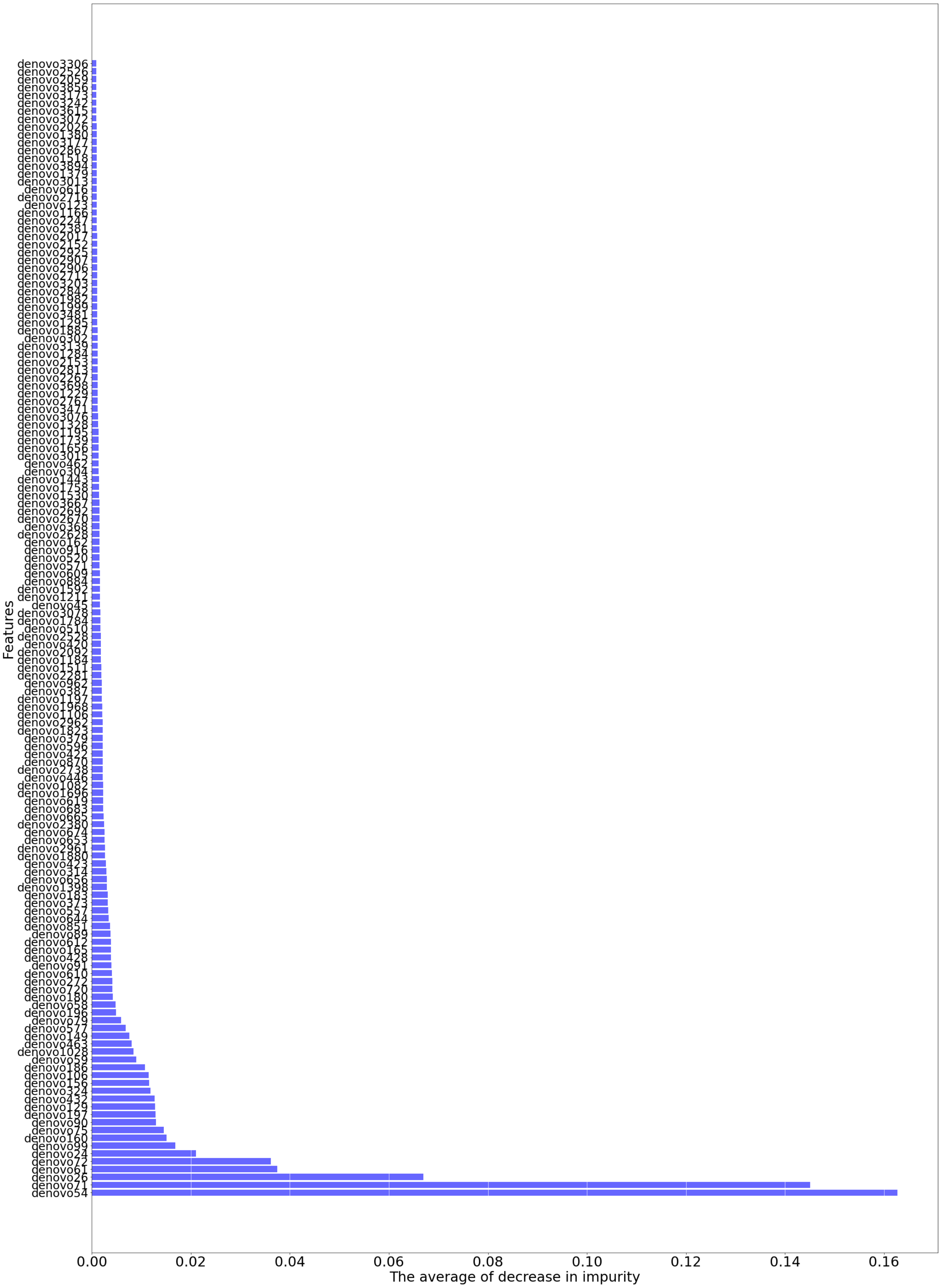


**Additional file 3: Figure S3**

Top 200 out of 3347 important features of the RF algorithm for the CDI case–control study.

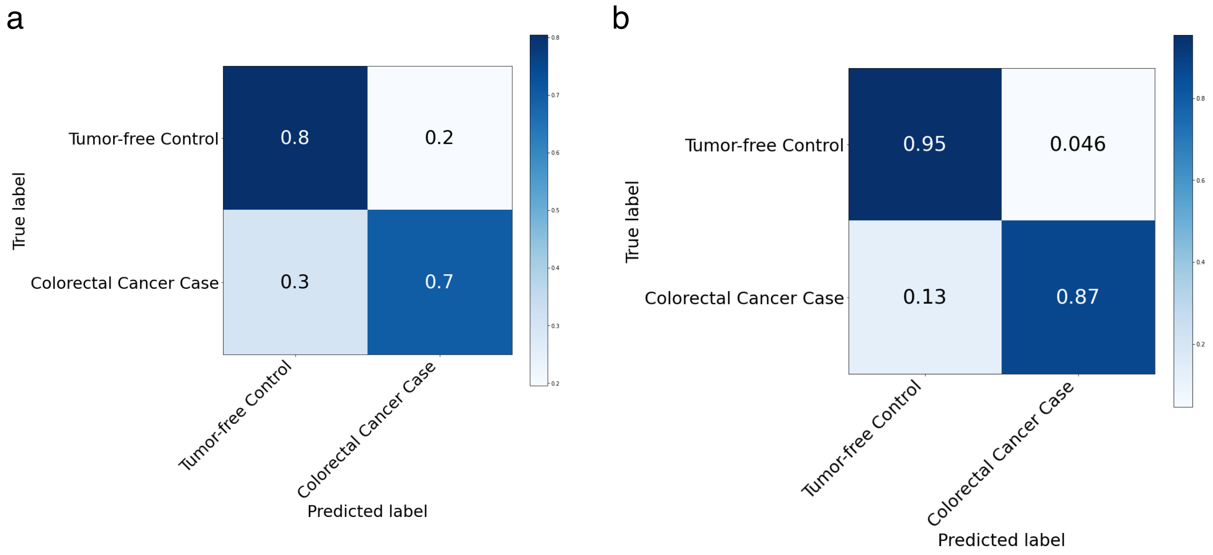


#### Additional file 4: Figure S4

Confusion matrices comparing the performance of random forest (with and without forward variable selection) for shotgun metagenomics data from the human colorectal cancer (CRC) dataset. **a** Random forest classifier for the shotgun metagenomics data; **b** Random forest classifier coupled with forward variable selection (RF-FVS) for the shotgun metagenomics data.

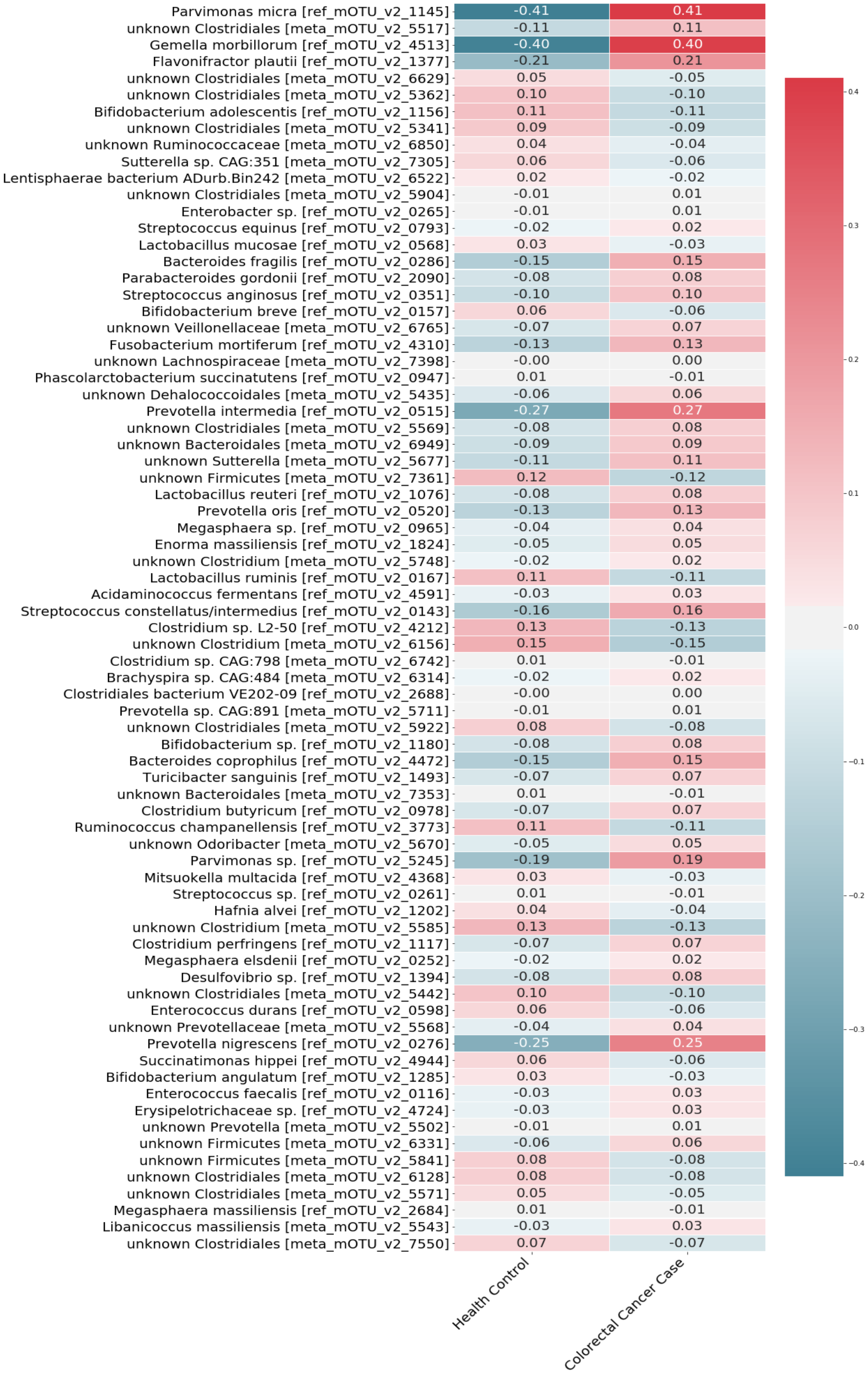


**Additional file 5: Figure S5** Correlation among the 75 selected species with colorectal cancer cases and tumor-free (healthy) controls.

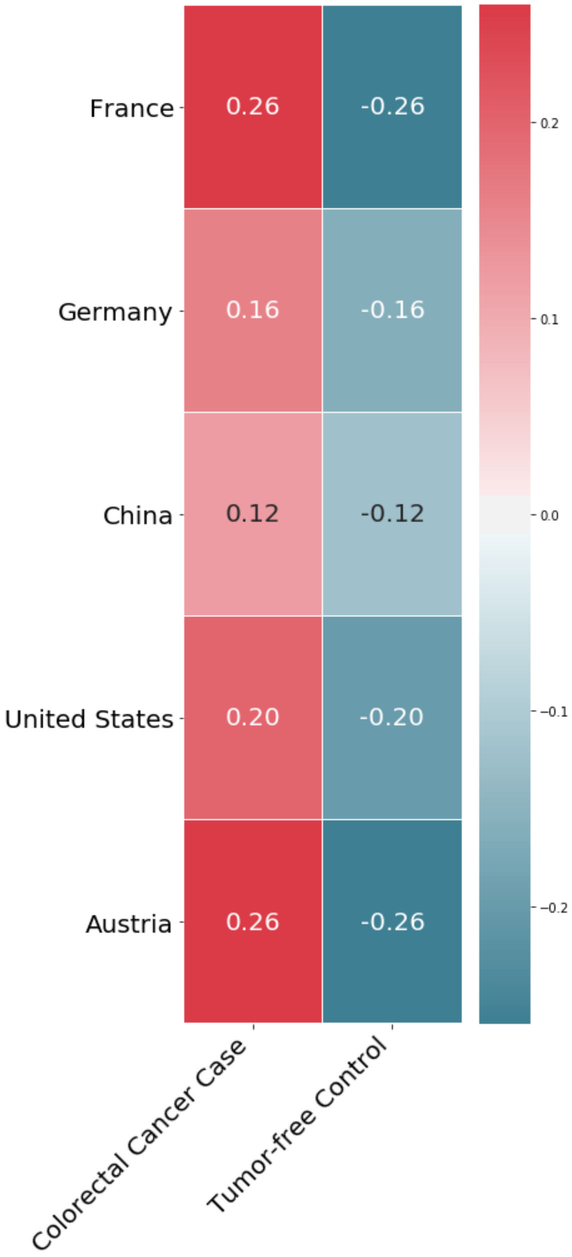


**Additional file 6: Figure S6**

Correlation of *Flavonifractor plautii* species with colorectal cancer cases and tumor-free controls of cohorts from France, Germany, China, United States, and Austria.

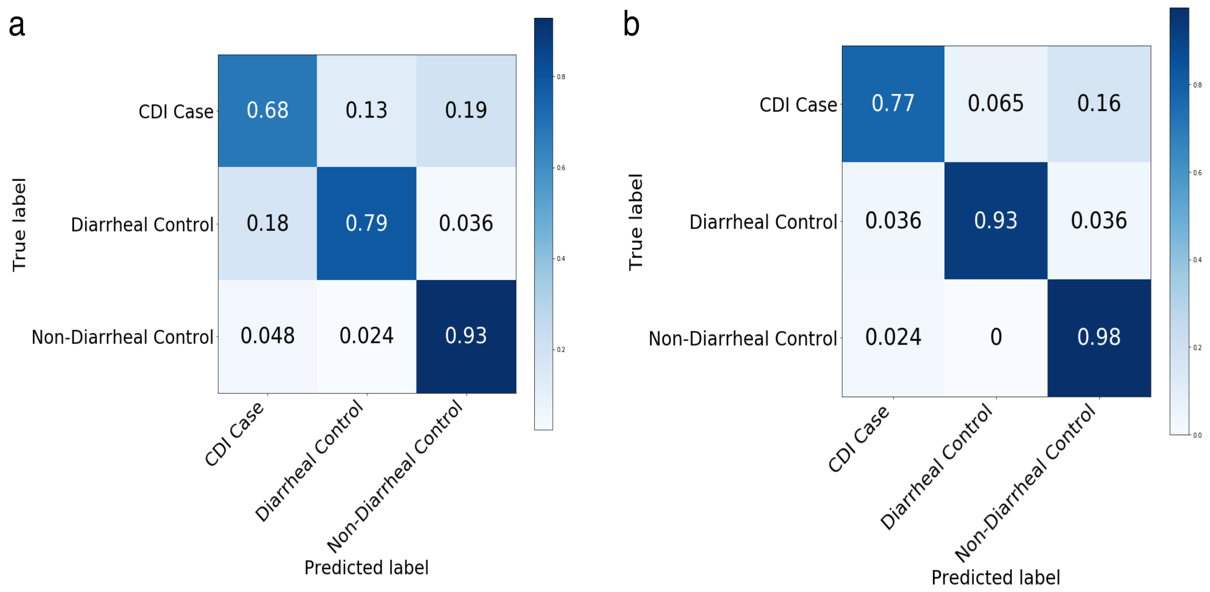


**Additional file 7: Figure S7**

Comparison of the performance of RF-FVS coupled with prescreen step for KEGG in the CDI data. **a** Random forest classifier for KEGG data; **b** Random forest classifier coupled with forward variable selection (RF-FVS) for KEGG data.

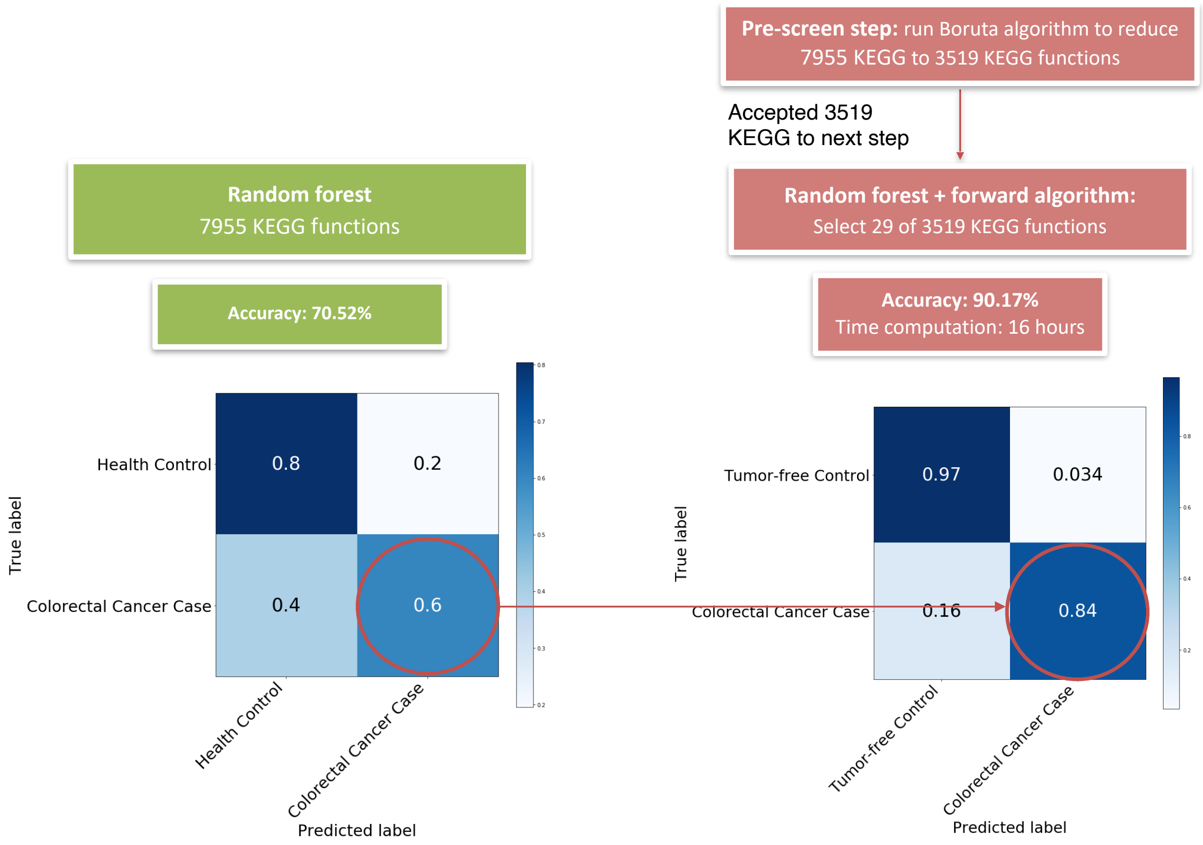


**Additional file 8: Figure S8**
Comparison of the performance of RF-FVS coupled with prescreen step for KEGG in the CRC shotgun metagenomic data.

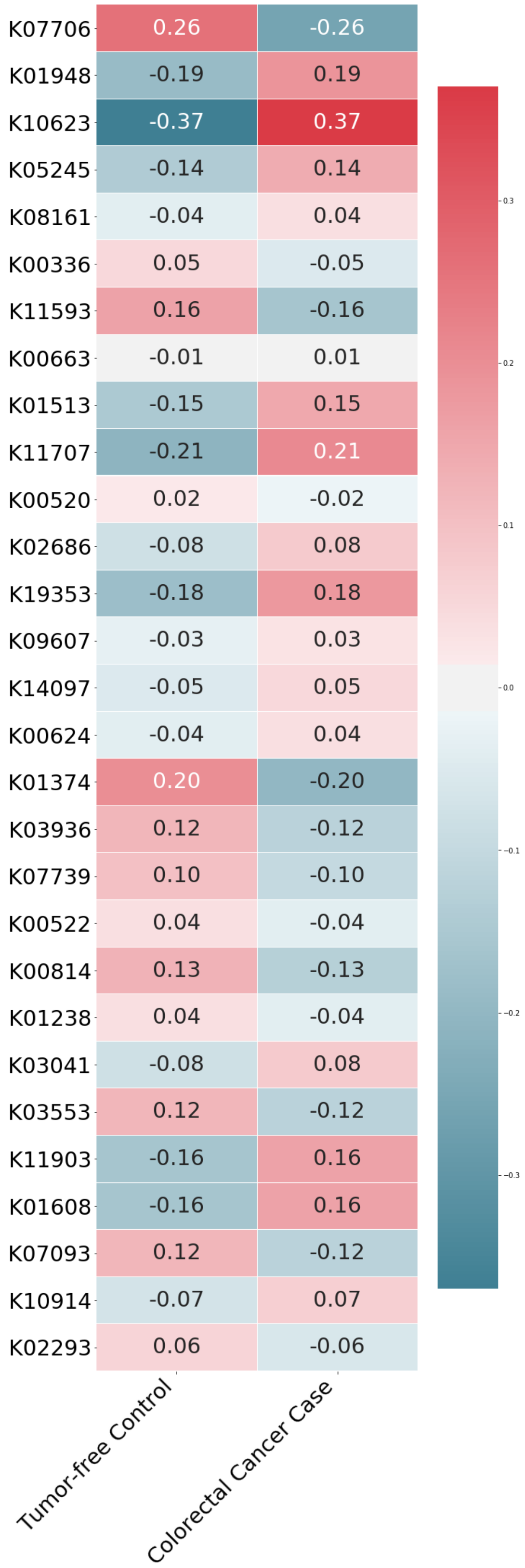


**Additional file 9: Figure S9**

Correlation among the 29 KEGG functions with colorectal cancer cases and tumor-free controls.

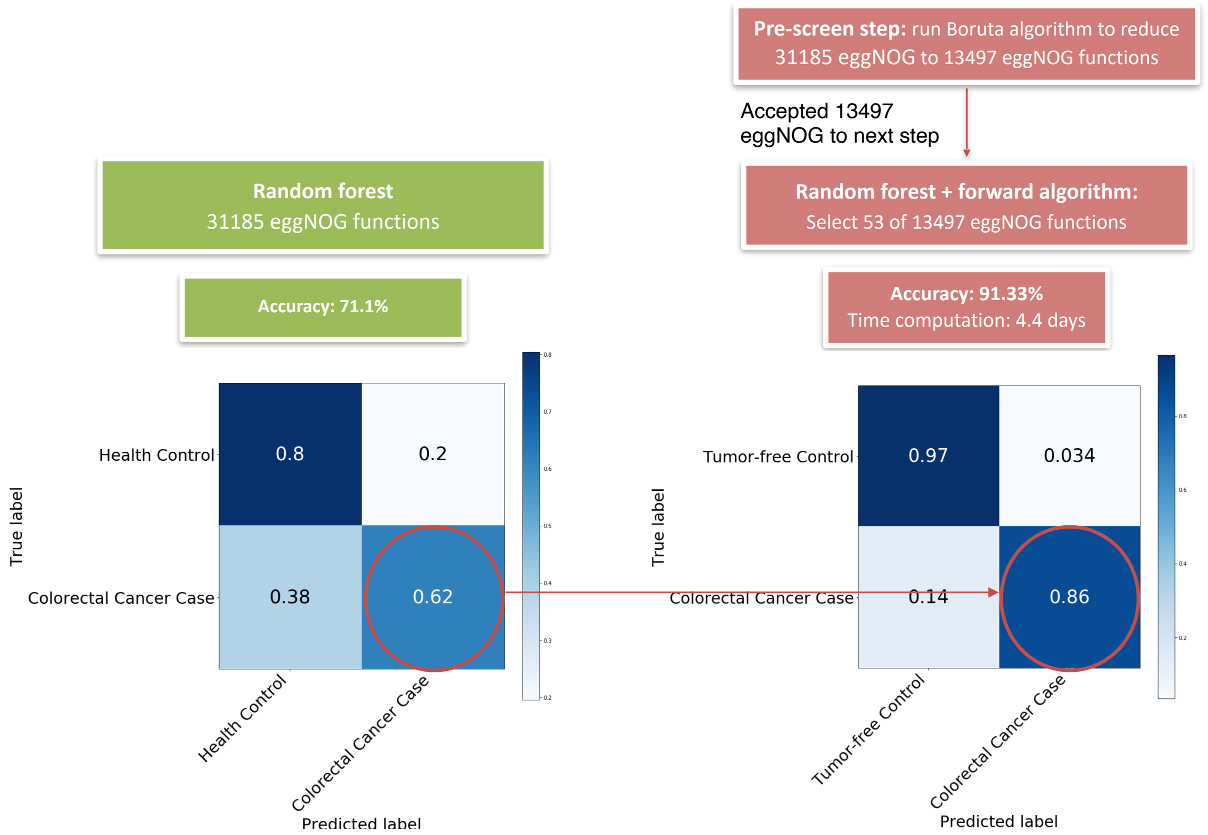


**Additional file 10: Figure S10**

Comparison of the performance of RF-FVS coupled with prescreen step for eggNOG in the CRC shotgun metagenomic data.

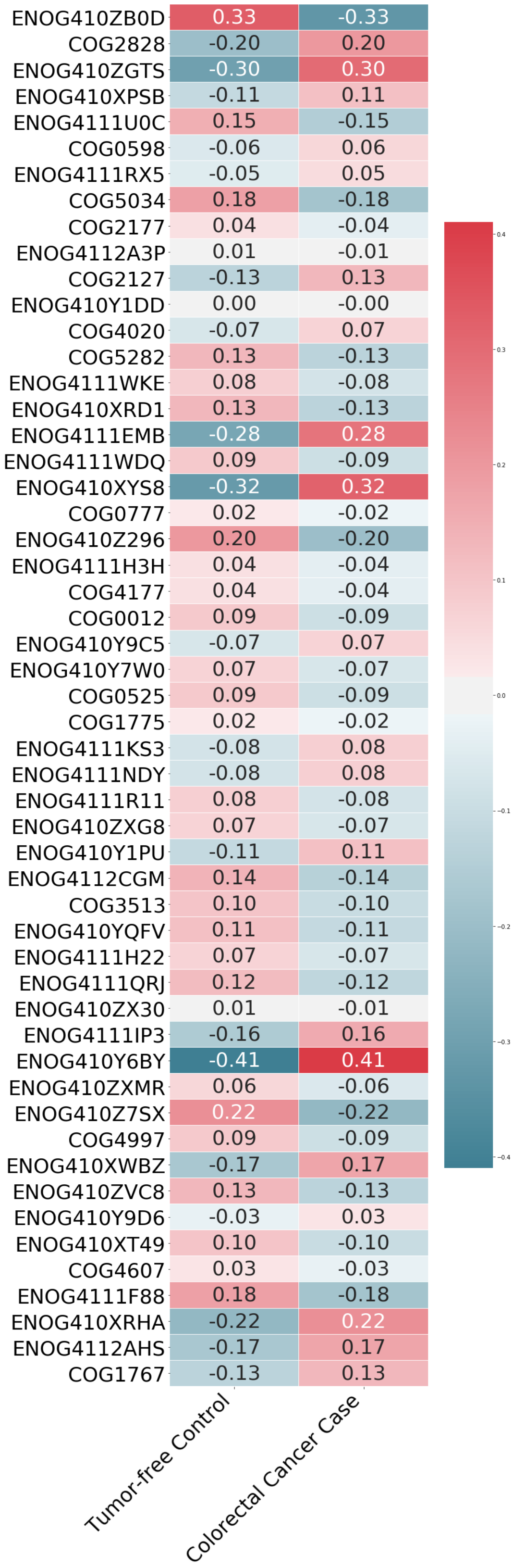


**Additional file 11: Figure S11**
Correlation among the 53 selected eggNOG functions with colorectal cancer cases and tumor-free controls.

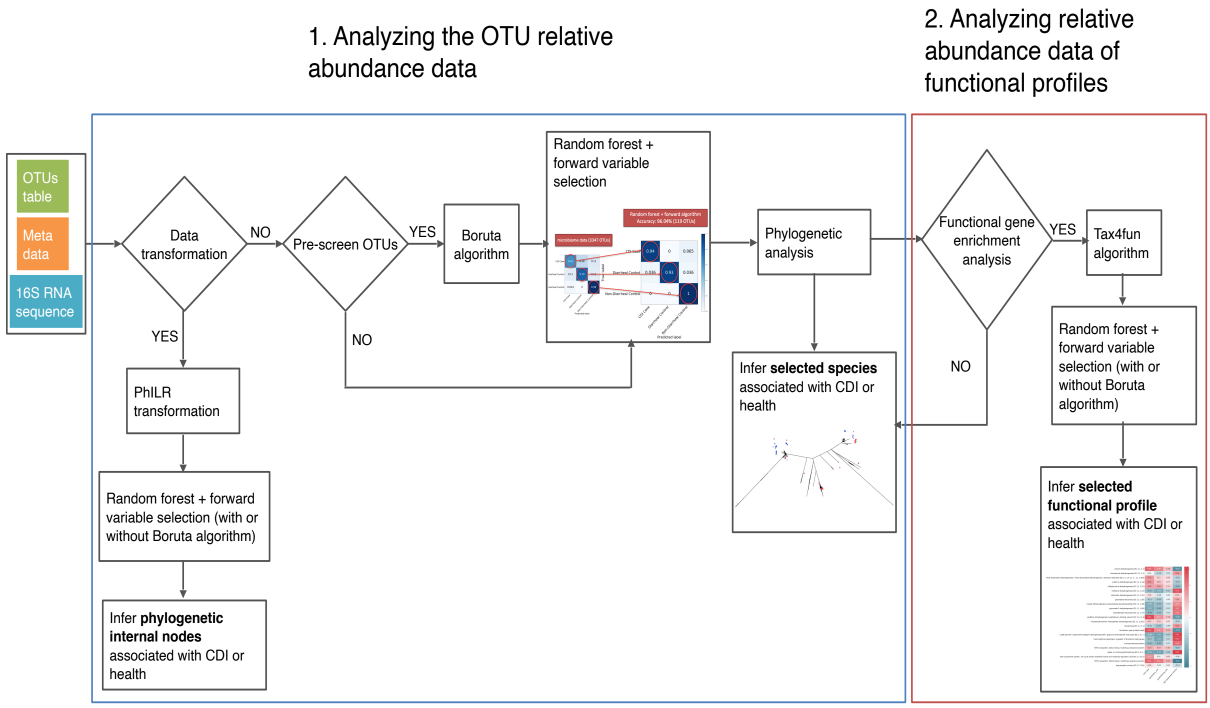


**Additional file 12: Figure S12**
Workflow for the new random forest coupled with forward variable selection (RF-FVS) approach developed and implemented in this study.

**Supplementary Table legends**

**Additional file 13: Table S1 a**

Measure of variable importance from the results of the random forest model for 3347 microbiome species in the CDI data.

| Species | Gini impurity | Species | Gini impurity | Species | Gini impurity |
|---|---|---|---|---|---|
| denovo54 | 0.1627 | denovo2019 | 0.0033 | ⋮ | ⋮ |
| denovo1238 | 0.1451 | denovo1773 | 0.0032 | | |
| denovo1326 | 0.0669 | denovo2528 | 0.0032 | | |
| denovo3072 | 0.0374 | denovo3205 | 0.0030 | denovo73 | 0.0002 |
| denovo601 | 0.0362 | denovo1888 | 0.0030 | denovo30 | 0.0002 |
| denovo163 | 0.0211 | denovo1776 | 0.0029 | denovo164 | 0.0002 |
| denovo3859 | 0.0169 | denovo1012 | 0.0029 | denovo40 | 0.0002 |
| denovo612 | 0.0151 | denovo2293 | 0.0027 | denovo1907 | 0.0002 |
| denovo2318 | 0.0146 | denovo1982 | 0.0026 | denovo428 | 0.0002 |
| denovo2346 | 0.0129 | denovo1485 | 0.0026 | denovo747 | 0.0002 |
| denovo3486 | 0.0129 | denovo2772 | 0.0026 | denovo349 | 0.0002 |
| denovo3031 | 0.0127 | denovo1775 | 0.0025 | denovo566 | 0.0002 |
| denovo1207 | 0.0127 | denovo302 | 0.0024 | denovo921 | 0.0002 |
| denovo3675 | 0.0118 | denovo1987 | 0.0023 | denovo653 | 0.0002 |
| denovo3583 | 0.0116 | denovo1517 | 0.0023 | denovo718 | 0.0002 |
| denovo3056 | 0.0114 | denovo2905 | 0.0023 | denovo1082 | 0.0002 |
| denovo423 | 0.0107 | denovo1190 | 0.0023 | denovo121 | 0.0002 |
| denovo162 | 0.0090 | denovo1251 | 0.0023 | denovo338 | 0.0002 |
| denovo156 | 0.0085 | denovo1539 | 0.0022 | denovo542 | 0.0002 |
| denovo1980 | 0.0080 | denovo1594 | 0.0022 | denovo672 | 0.0002 |
| denovo3851 | 0.0076 | denovo1123 | 0.0022 | denovo1072 | 0.0002 |
| denovo2341 | 0.0068 | denovo1774 | 0.0022 | denovo475 | 0.0002 |
| denovo2508 | 0.0059 | denovo3059 | 0.0022 | denovo885 | 0.0002 |
| denovo3050 | 0.0049 | denovo1988 | 0.0022 | denovo1792 | 0.0002 |
| denovo2479 | 0.0048 | denovo3058 | 0.0022 | denovo661 | 0.0002 |
| denovo2189 | 0.0042 | denovo3053 | 0.0022 | denovo2117 | 0.0002 |
| denovo127 | 0.0042 | denovo1983 | 0.0021 | denovo765 | 0.0002 |
| denovo3365 | 0.0041 | denovo1106 | 0.0021 | denovo280 | 0.0002 |
| denovo788 | 0.0041 | denovo1981 | 0.0020 | denovo614 | 0.0001 |
| denovo1089 | 0.0039 | denovo618 | 0.0020 | denovo521 | 0.0000 |
| denovo1399 | 0.0039 | denovo3698 | 0.0020 | denovo1999 | 0.0000 |
| denovo1519 | 0.0038 | denovo3204 | 0.0019 | denovo1649 | 0.0000 |

| | | | | | |
|---|---|---|---|---|---|
| denovo617 | 0.0038 | denovo1278 | 0.0019 | denovo1000 | 0.0000 |
| denovo3207 | 0.0038 | denovo610 | 0.0018 | denovo73 | 0.0000 |
| denovo695 | 0.0037 | denovo1370 | 0.0018 | denovo30 | 0.0000 |
| denovo26 | 0.0034 | denovo3201 | 0.0018 | denovo164 | 0.0000 |

**Additional file 13: Table S1 b**

Selected species in three groups associated with CDI case and diarrheal and non-diarrheal controls. Information about family and genus are shown. Gray shading indicates species that are associated with CDI disease.

| | Species ID | Over-representation in | | |
|---|---|---|---|---|
| | | CDI case | Non-diarrheal control | Diarrheal control |
| Group A | denovo2019 | ✓ | × | × |
| | denovo1773 | ✓ | × | × |
| | denovo2528 | ✓ | × | ✓ |
| | denovo3205 | ✓ | × | ✓ |
| | denovo1888 | × | ✓ | × |
| | denovo1776 | × | ✓ | × |
| | denovo1012 | × | ✓ | × |
| | denovo2293 | × | ✓ | × |
| | denovo1982 | × | ✓ | × |
| | denovo1485 | × | ✓ | × |
| | denovo2772 | × | ✓ | × |
| | denovo1775 | × | ✓ | × |
| | denovo302 | × | ✓ | × |
| | denovo1987 | × | ✓ | × |
| | denovo1517 | × | ✓ | × |
| | denovo2905 | × | ✓ | × |
| | denovo1190 | × | ✓ | × |
| | denovo1251 | × | ✓ | × |
| | denovo1539 | × | ✓ | × |
| | denovo1594 | × | ✓ | × |
| | denovo1123 | × | ✓ | × |
| | denovo1774 | × | ✓ | × |
| | denovo3059 | × | ✓ | × |
| | denovo1988 | × | ✓ | × |
| | denovo3058 | × | ✓ | × |
| | denovo3053 | × | ✓ | × |
| | denovo1983 | × | ✓ | × |
| | denovo1106 | × | ✓ | × |

| | | | | |
|---|---|---|---|---|
| | denovo1981 | × | ✓ | × |
| | denovo618 | × | ✓ | × |
| | denovo3698 | × | ✓ | × |
| | denovo3204 | × | ✓ | × |
| | denovo1278 | × | ✓ | × |
| | denovo610 | × | ✓ | × |
| | denovo1370 | × | ✓ | × |
| | denovo3201 | × | ✓ | × |
| | denovo3057 | × | ✓ | × |
| | denovo1187 | × | ✓ | × |
| | denovo2923 | × | ✓ | × |
| | denovo2888 | × | ✓ | × |
| | denovo3692 | × | ✓ | × |
| | denovo1778 | × | ✓ | × |
| | denovo615 | × | ✓ | × |
| | denovo1510 | × | ✓ | × |
| | denovo613 | × | ✓ | × |
| | denovo1105 | × | ✓ | × |
| | denovo3203 | × | ✓ | × |
| | denovo3054 | × | ✓ | × |
| | denovo611 | × | ✓ | × |
| | denovo3630 | × | ✓ | × |
| | denovo557 | × | ✓ | × |
| | denovo2268 | × | ✓ | × |
| | denovo2526 | × | ✓ | × |
| | denovo679 | × | ✓ | × |
| **Group B** | denovo54 | ✓ | × | × |
| | denovo1238 | ✓ | × | ✓ |
| | denovo1326 | ✓ | × | × |
| | denovo3072 | ✓ | × | ✓ |
| | denovo601 | ✓ | × | ✓ |
| | denovo163 | ✓ | × | ✓ |
| | denovo3859 | ✓ | × | × |
| | denovo612 | ✓ | × | ✓ |
| | denovo2318 | ✓ | × | ✓ |
| | denovo2346 | ✓ | × | ✓ |
| | denovo3486 | ✓ | × | ✓ |
| | denovo3031 | ✓ | × | ✓ |
| | denovo1207 | ✓ | × | ✓ |
| | denovo3675 | ✓ | × | × |
| | denovo3583 | ✓ | × | ✓ |
| | denovo3056 | × | ✓ | × |
| | denovo423 | × | ✓ | ✓ |
| | denovo162 | × | ✓ | × |
| | denovo156 | × | ✓ | × |
| **Group C** | denovo1980 | ✓ | × | ✓ |
| | denovo3851 | ✓ | × | ✓ |
| | denovo2341 | ✓ | × | ✓ |

| | | | |
|---|---|---|---|
| denovo2508 | ✓ | ✗ | ✓ |
| denovo3050 | ✓ | ✗ | ✗ |
| denovo2479 | ✓ | ✗ | ✓ |
| denovo2189 | ✓ | ✗ | ✗ |
| denovo127 | ✓ | ✗ | ✗ |
| denovo3365 | ✓ | ✗ | ✗ |
| denovo788 | ✓ | ✗ | ✓ |
| denovo1089 | ✓ | ✗ | ✓ |
| denovo1399 | ✓ | ✗ | ✓ |
| denovo1519 | ✓ | ✗ | ✗ |
| denovo617 | ✓ | ✗ | ✗ |
| denovo3207 | ✓ | ✗ | ✓ |
| denovo695 | ✓ | ✗ | ✓ |
| denovo26 | ✗ | ✓ | |
| denovo1752 | ✗ | ✓ | ✗ |
| denovo9 | ✗ | ✓ | ✗ |
| denovo3274 | ✗ | ✓ | ✗ |
| denovo2467 | ✗ | ✓ | ✗ |
| denovo3206 | ✗ | ✓ | ✗ |
| denovo374 | ✗ | ✓ | ✗ |
| denovo1295 | ✗ | ✓ | ✗ |
| denovo3269 | ✗ | ✓ | ✗ |
| denovo1777 | ✗ | ✓ | ✗ |
| denovo447 | ✗ | ✓ | ✗ |
| denovo1354 | ✗ | ✓ | ✓ |
| denovo2159 | ✗ | ✓ | ✗ |
| denovo1772 | ✗ | ✓ | ✗ |
| denovo347 | ✗ | ✓ | ✗ |
| denovo1756 | ✗ | ✓ | ✗ |
| denovo3055 | ✗ | ✓ | ✗ |
| denovo1779 | ✗ | ✓ | ✗ |
| denovo2753 | ✗ | ✓ | ✗ |
| denovo1999 | ✗ | ✓ | ✗ |

**Additional file 13: Table S1 c**

Taxonomy information for the species listed. Gray shading indicates species that are associated with CDI disease.

| | Species ID | Taxonomy |
|---|---|---|
| Group A | denovo2019 | k__Bacteria;p__Bacteroidetes;c__Bacteroidia;o__Bacteroidales;f__Bacteroidaceae;g__Bacteroides;s__;d__denovo2019 |
| | denovo1773 | k__Bacteria;p__Firmicutes;c__Clostridia;o__Clostridiales;f__Lachnospiraceae;g__Clostridium_XlVb;s__;d__denovo1773 |
| | denovo2528 | k__Bacteria;p__Bacteroidetes;c__Bacteroidia;o__Bacteroidales;f__Rikenellaceae;g__Alistipes;s__;d__denovo2528 |

| | | |
|---|---|---|
| | denovo3205 | k__Bacteria;p__Bacteroidetes;c__Bacteroidia;o__Bacteroidales;f__Porphyromonadaceae;g__Parabacteroides;s__;d__denovo3205 |
| | denovo1888 | k__Bacteria;p__Bacteroidetes;c__Bacteroidia;o__Bacteroidales;f__Bacteroidaceae;g__Bacteroides;s__;d__denovo1888 |
| | denovo1776 | k__Bacteria;p__Bacteroidetes;c__Bacteroidia;o__Bacteroidales;f__Porphyromonadaceae;g__Parabacteroides;s__;d__denovo1776 |
| | denovo1012 | k__Bacteria;p__Bacteroidetes;c__Bacteroidia;o__Bacteroidales;f__Bacteroidaceae;g__Bacteroides;s__;d__denovo1012 |
| | denovo2293 | k__Bacteria;p__Bacteroidetes;c__Bacteroidia;o__Bacteroidales;f__Porphyromonadaceae;g__Barnesiella;s__;d__denovo2293 |
| | denovo1982 | k__Bacteria;p__Bacteroidetes;c__Bacteroidia;o__Bacteroidales;f__Bacteroidaceae;g__Bacteroides;s__;d__denovo1982 |
| | denovo1485 | k__Bacteria;p__Bacteroidetes;c__Bacteroidia;o__Bacteroidales;f__Bacteroidaceae;g__Bacteroides;s__;d__denovo1485 |
| | denovo2772 | k__Bacteria;p__Firmicutes;c__Erysipelotrichia;o__Erysipelotrichales;f__Erysipelotrichaceae;g__Clostridium_XVIII;s__;d__denovo2772 |
| | denovo1775 | k__Bacteria;p__Bacteroidetes;c__Bacteroidia;o__Bacteroidales;f__Bacteroidaceae;g__Bacteroides;s__;d__denovo1775 |
| | denovo302 | k__Bacteria;p__Bacteroidetes;c__Bacteroidia;o__Bacteroidales;f__Rikenellaceae;g__Alistipes;s__;d__denovo302 |
| | denovo1987 | k__Bacteria;p__Bacteroidetes;c__Bacteroidia;o__Bacteroidales;f__Bacteroidaceae;g__Bacteroides;s__;d__denovo1987 |
| | denovo1517 | k__Bacteria;p__Bacteroidetes;c__Bacteroidia;o__Bacteroidales;f__Porphyromonadaceae;g__Barnesiella;s__;d__denovo1517 |
| | denovo2905 | k__Bacteria;p__Bacteroidetes;c__Bacteroidia;o__Bacteroidales;f__Bacteroidaceae;g__Bacteroides;s__;d__denovo2905 |
| | denovo1190 | k__Bacteria;p__Bacteroidetes;c__Bacteroidia;o__Bacteroidales;f__Porphyromonadaceae;g__Parabacteroides;s__;d__denovo1190 |
| | denovo1251 | k__Bacteria;p__Bacteroidetes;c__Bacteroidia;o__Bacteroidales;f__Bacteroidaceae;g__Bacteroides;s__;d__denovo1251 |
| | denovo1539 | k__Bacteria;p__Bacteroidetes;c__Bacteroidia;o__Bacteroidales;f__Rikenellaceae;g__Alistipes;s__;d__denovo1539 |
| | denovo1594 | k__Bacteria;p__Bacteroidetes;c__Bacteroidia;o__Bacteroidales;f__Bacteroidaceae;g__Bacteroides;s__;d__denovo1594 |
| | denovo1123 | k__Bacteria;p__Bacteroidetes;c__Bacteroidia;o__Bacteroidales;f__Bacteroidaceae;g__Bacteroides;s__;d__denovo1123 |

| | | |
|---|---|---|
| | denovo1774 | k__Bacteria;p__Bacteroidetes;c__Bacteroidia;o__Bacteroidales;f__Bacteroidaceae;g__Bacteroides;s__;d__denovo1774 |
| | denovo3059 | k__Bacteria;p__Bacteroidetes;c__Bacteroidia;o__Bacteroidales;f__Bacteroidaceae;g__Bacteroides;s__;d__denovo3059 |
| | denovo1988 | k__Bacteria;p__Bacteroidetes;c__Bacteroidia;o__Bacteroidales;f__Bacteroidaceae;g__Bacteroides;s__;d__denovo1988 |
| | denovo3058 | k__Bacteria;p__Bacteroidetes;c__Bacteroidia;o__Bacteroidales;f__Bacteroidaceae;g__Bacteroides;s__;d__denovo3058 |
| | denovo3053 | k__Bacteria;p__Bacteroidetes;c__Bacteroidia;o__Bacteroidales;f__Bacteroidaceae;g__Bacteroides;s__;d__denovo3053 |
| | denovo1983 | k__Bacteria;p__Bacteroidetes;c__Bacteroidia;o__Bacteroidales;f__Bacteroidaceae;g__Bacteroides;s__;d__denovo1983 |
| | denovo1106 | k__Bacteria;p__Bacteroidetes;c__Bacteroidia;o__Bacteroidales;f__Porphyromonadaceae;g__Parabacteroides;s__;d__denovo1106 |
| | denovo1981 | k__Bacteria;p__Bacteroidetes;c__Bacteroidia;o__Bacteroidales;f__Bacteroidaceae;g__Bacteroides;s__;d__denovo1981 |
| | denovo618 | k__Bacteria;p__Bacteroidetes;c__Bacteroidia;o__Bacteroidales;f__Bacteroidaceae;g__Bacteroides;s__;d__denovo618 |
| | denovo3698 | k__Bacteria;p__Bacteroidetes;c__Bacteroidia;o__Bacteroidales;f__Porphyromonadaceae;g__Parabacteroides;s__;d__denovo3698 |
| | denovo3204 | k__Bacteria;p__Bacteroidetes;c__Bacteroidia;o__Bacteroidales;f__Bacteroidaceae;g__Bacteroides;s__;d__denovo3204 |
| | denovo1278 | k__Bacteria;p__Bacteroidetes;c__Bacteroidia;o__Bacteroidales;f__Bacteroidaceae;g__Bacteroides;s__;d__denovo1278 |
| | denovo610 | k__Bacteria;p__Bacteroidetes;c__Bacteroidia;o__Bacteroidales;f__Bacteroidaceae;g__Bacteroides;s__;d__denovo610 |
| | denovo1370 | k__Bacteria;p__Bacteroidetes;c__Bacteroidia;o__Bacteroidales;f__Porphyromonadaceae;g__Parabacteroides;s__;d__denovo1370 |
| | denovo3201 | k__Bacteria;p__Bacteroidetes;c__Bacteroidia;o__Bacteroidales;f__Bacteroidaceae;g__Bacteroides;s__;d__denovo3201 |
| | denovo3057 | k__Bacteria;p__Bacteroidetes;c__Bacteroidia;o__Bacteroidales;f__Porphyromonadaceae;g__Barnesiella;s__;d__denovo3057 |
| | denovo1187 | k__Bacteria;p__Bacteroidetes;c__Bacteroidia;o__Bacteroidales;f__Porphyromonadaceae;g__Parabacteroides;s__;d__denovo1187 |
| | denovo2923 | k__Bacteria;p__Bacteroidetes;c__Bacteroidia;o__Bacteroidales;f__Prevotellaceae;g__Xylanibacter;s__;d__denovo2923 |

| | | |
|---|---|---|
| | denovo2888 | k__Bacteria;p__Bacteroidetes;c__Bacteroidia;o__Bacteroidales;f__Bacteroidaceae;g__Bacteroides;s__;d__denovo2888 |
| | denovo3692 | k__Bacteria;p__Bacteroidetes;c__Bacteroidia;o__Bacteroidales;f__Bacteroidaceae;g__Bacteroides;s__;d__denovo3692 |
| | denovo1778 | k__Bacteria;p__Bacteroidetes;c__Bacteroidia;o__Bacteroidales;f__Bacteroidaceae;g__Bacteroides;s__;d__denovo1778 |
| | denovo615 | k__Bacteria;p__Bacteroidetes;c__Bacteroidia;o__Bacteroidales;f__Bacteroidaceae;g__Bacteroides;s__;d__denovo615 |
| | denovo1510 | k__Bacteria;p__Bacteroidetes;c__Bacteroidia;o__Bacteroidales;f__Bacteroidaceae;g__Bacteroides;s__;d__denovo1510 |
| | denovo613 | k__Bacteria;p__Bacteroidetes;c__Bacteroidia;o__Bacteroidales;f__Porphyromonadaceae;g__Parabacteroides;s__;d__denovo613 |
| | denovo1105 | k__Bacteria;p__Bacteroidetes;c__Bacteroidia;o__Bacteroidales;f__Bacteroidaceae;g__Bacteroides;s__;d__denovo1105 |
| | denovo3203 | k__Bacteria;p__Bacteroidetes;c__Bacteroidia;o__Bacteroidales;f__Porphyromonadaceae;g__Parabacteroides;s__;d__denovo3203 |
| | denovo3054 | k__Bacteria;p__Firmicutes;c__Clostridia;o__Clostridiales;f__Ruminococcaceae;g__Hydrogenoanaerobacterium;s__;d__denovo3054 |
| | denovo611 | k__Bacteria;p__Bacteroidetes;c__Bacteroidia;o__Bacteroidales;f__Bacteroidaceae;g__Bacteroides;s__;d__denovo611 |
| | denovo3630 | k__Bacteria;p__Bacteroidetes;c__Bacteroidia;o__Bacteroidales;f__Porphyromonadaceae;g__Barnesiella;s__;d__denovo3630 |
| | denovo557 | k__Bacteria;p__Bacteroidetes;c__Bacteroidia;o__Bacteroidales;f__Bacteroidaceae;g__Bacteroides;s__;d__denovo557 |
| | denovo2268 | k__Bacteria;p__Bacteroidetes;c__Bacteroidia;o__Bacteroidales;f__Bacteroidaceae;g__Bacteroides;s__;d__denovo2268 |
| | denovo2526 | k__Bacteria;p__Bacteroidetes;c__Bacteroidia;o__Bacteroidales;f__Bacteroidaceae;g__Bacteroides;s__;d__denovo2526 |
| | denovo679 | k__Bacteria;p__Bacteroidetes;c__Bacteroidia;o__Bacteroidales;f__Porphyromonadaceae;g__Parabacteroides;s__;d__denovo679 |
| **Group B** | denovo54 | k__Bacteria;p__Firmicutes;c__Clostridia;o__Clostridiales;f__Peptostreptococcaceae;g__Clostridium_XI;s__;d__denovo54 |
| | denovo1238 | k__Bacteria;p__Verrucomicrobia;c__Verrucomicrobiae;o__Verrucomicrobiales;f__Verrucomicrobiaceae;g__Akkermansia;s__;d__denovo1238 |
| | denovo1326 | k__Bacteria;p__Firmicutes;c__Bacilli;o__Lactobacillales;f__Lactobacillaceae;g__Lactobacillus;s__;d__denovo1326 |

| | | |
|---|---|---|
| | denovo3072 | k__Bacteria;p__Firmicutes;c__Bacilli;o__Lactobacillales;f__Enterococcaceae;g__Enterococcus;s__;d__denovo3072 |
| | denovo601 | k__Bacteria;p__Actinobacteria;c__Actinobacteria;o__Coriobacteriales;f__Coriobacteriaceae;g__Eggerthella;s__;d__denovo601 |
| | denovo163 | k__Bacteria;p__Firmicutes;c__Clostridia;o__Clostridiales;f__Eubacteriaceae;g__;s__;d__denovo163 |
| | denovo3859 | k__Bacteria;p__Verrucomicrobia;c__Verrucomicrobiae;o__Verrucomicrobiales;f__Verrucomicrobiaceae;g__Akkermansia;s__;d__denovo3859 |
| | denovo612 | k__Bacteria;p__Firmicutes;c__Bacilli;o__Lactobacillales;f__Enterococcaceae;g__Enterococcus;s__;d__denovo612 |
| | denovo2318 | k__Bacteria;p__Verrucomicrobia;c__Verrucomicrobiae;o__Verrucomicrobiales;f__Verrucomicrobiaceae;g__Akkermansia;s__;d__denovo2318 |
| | denovo2346 | k__Bacteria;p__Firmicutes;c__Negativicutes;o__Selenomonadales;f__Veillonellaceae;g__Veillonella;s__;d__denovo2346 |
| | denovo3486 | k__Bacteria;p__Firmicutes;c__Negativicutes;o__Selenomonadales;f__Veillonellaceae;g__Veillonella;s__;d__denovo3486 |
| | denovo3031 | k__Bacteria;p__Firmicutes;c__Bacilli;o__Lactobacillales;f__Enterococcaceae;g__Enterococcus;s__;d__denovo3031 |
| | denovo1207 | k__Bacteria;p__Firmicutes;c__Bacilli;o__Lactobacillales;f__Enterococcaceae;g__Enterococcus;s__;d__denovo1207 |
| | denovo3675 | k__Bacteria;p__Firmicutes;c__Negativicutes;o__Selenomonadales;f__Veillonellaceae;g__Dialister;s__;d__denovo3675 |
| | denovo3583 | k__Bacteria;p__Firmicutes;c__Bacilli;o__Lactobacillales;f__Lactobacillaceae;g__Lactobacillus;s__;d__denovo3583 |
| | denovo3056 | k__Bacteria;p__Firmicutes;c__Negativicutes;o__Selenomonadales;f__;g__;s__;d__denovo3056 |
| | denovo423 | k__Bacteria;p__Firmicutes;c__Clostridia;o__Clostridiales;f__;g__;s__;d__denovo423 |
| | denovo162 | k__Bacteria;p__Firmicutes;c__Clostridia;o__Clostridiales;f__;g__;s__;d__denovo162 |
| | denovo156 | k__Bacteria;p__Firmicutes;c__Clostridia;o__Clostridiales;f__Ruminococcaceae;g__;s__;d__denovo156 |
| **Group C** | denovo1980 | k__Bacteria;p__Proteobacteria;c__Gammaproteobacteria;o__Enterobacteriales;f__Enterobacteriaceae;g__;s__;d__denovo1980 |
| | denovo3851 | k__Bacteria;p__Proteobacteria;c__Gammaproteobacteria;o__;f__;g__;s__;d__denovo3851 |
| | denovo2341 | k__Bacteria;p__Proteobacteria;c__Gammaproteobacteria;o__Enterobacteriales;f__Enterobacteriaceae;g__;s__;d__denovo2341 |
| | denovo2508 | k__Bacteria;p__Proteobacteria;c__Gammaproteobacteria;o__Enterobacteriales;f__Enterobacteriaceae;g__;s__;d__denovo2508 |
| | denovo3050 | k__Bacteria;p__Firmicutes;c__Clostridia;o__Clostridiales;f__Lachnospiraceae;g__Blautia;s__;d__denovo3050 |
| | denovo2479 | k__Bacteria;p__Proteobacteria;c__Gammaproteobacteria;o__Enterobacteriales;f__Enterobacteriaceae;g__;s__;d__denovo2479 |

| | | |
|---|---|---|
| | denovo2189 | k__Bacteria;p__Proteobacteria;c__Gammaproteobacteria;o__Enterobacteriales;f__Enterobacteriaceae;g__Raoultella;s__;d__denovo2189 |
| | denovo127 | k__Bacteria;p__Proteobacteria;c__Gammaproteobacteria;o__Enterobacteriales;f__Enterobacteriaceae;g__Proteus;s__;d__denovo127 |
| | denovo3365 | k__Bacteria;p__Firmicutes;c__Clostridia;o__Clostridiales;f__Lachnospiraceae;g__Ruminococcus2;s__;d__denovo3365 |
| | denovo788 | k__Bacteria;p__Proteobacteria;c__Gammaproteobacteria;o__Enterobacteriales;f__Enterobacteriaceae;g__;s__;d__denovo788 |
| | denovo1089 | k__Bacteria;p__Proteobacteria;c__Gammaproteobacteria;o__Enterobacteriales;f__Enterobacteriaceae;g__;s__;d__denovo1089 |
| | denovo1399 | k__Bacteria;p__Firmicutes;c__Clostridia;o__Clostridiales;f__Lachnospiraceae;g__Blautia;s__;d__denovo1399 |
| | denovo1519 | k__Bacteria;p__Firmicutes;c__Clostridia;o__Clostridiales;f__Lachnospiraceae;g__Blautia;s__;d__denovo1519 |
| | denovo617 | k__Bacteria;p__Firmicutes;c__Clostridia;o__Clostridiales;f__Lachnospiraceae;g__Robinsoniella;s__;d__denovo617 |
| | denovo3207 | k__Bacteria;p__Firmicutes;c__Clostridia;o__Clostridiales;f__Lachnospiraceae;g__Blautia;s__;d__denovo3207 |
| | denovo695 | k__Bacteria;p__Proteobacteria;c__Gammaproteobacteria;o__Enterobacteriales;f__Enterobacteriaceae;g__;s__;d__denovo695 |
| | denovo26 | k__Bacteria;p__Firmicutes;c__Clostridia;o__Clostridiales;f__Lachnospiraceae;g__;s__;d__denovo26 |
| | denovo1752 | k__Bacteria;p__Firmicutes;c__Clostridia;o__Clostridiales;f__Lachnospiraceae;g__;s__;d__denovo1752 |
| | denovo9 | k__Bacteria;p__Firmicutes;c__Clostridia;o__Clostridiales;f__Ruminococcaceae;g__Faecalibacterium;s__;d__denovo9 |
| | denovo3274 | k__Bacteria;p__Firmicutes;c__Clostridia;o__Clostridiales;f__Lachnospiraceae;g__;s__;d__denovo3274 |
| | denovo2467 | k__Bacteria;p__Proteobacteria;c__Gammaproteobacteria;o__Enterobacteriales;f__Enterobacteriaceae;g__;s__;d__denovo2467 |
| | denovo3206 | k__Bacteria;p__Firmicutes;c__Clostridia;o__Clostridiales;f__Lachnospiraceae;g__Blautia;s__;d__denovo3206 |
| | denovo374 | k__Bacteria;p__Firmicutes;c__Negativicutes;o__Selenomonadales;f__Veillonellaceae;g__;s__;d__denovo374 |
| | denovo1295 | k__Bacteria;p__Firmicutes;c__Clostridia;o__Clostridiales;f__Lachnospiraceae;g__Dorea;s__;d__denovo1295 |
| | denovo3269 | k__Bacteria;p__Firmicutes;c__Clostridia;o__Clostridiales;f__Lachnospiraceae;g__Coprococcus;s__;d__denovo3269 |
| | denovo1777 | k__Bacteria;p__Firmicutes;c__Clostridia;o__Clostridiales;f__Lachnospiraceae;g__Blautia;s__;d__denovo1777 |
| | denovo447 | k__Bacteria;p__Firmicutes;c__Clostridia;o__Clostridiales;f__Lachnospiraceae;g__Blautia;s__;d__denovo447 |
| | denovo1354 | k__Bacteria;p__Firmicutes;c__Clostridia;o__Clostridiales;f__Lachnospiraceae;g__Clostridium_XlVa;s__;d__denovo1354 |
| | denovo2159 | k__Bacteria;p__Firmicutes;c__Clostridia;o__Clostridiales;f__Ruminococcaceae;g__;s__;d__denovo2159 |
| | denovo1772 | k__Bacteria;p__Firmicutes;c__Clostridia;o__Clostridiales;f__Lachnospiraceae;g__Blautia;s__;d__denovo1772 |
| | denovo347 | k__Bacteria;p__Firmicutes;c__Clostridia;o__Clostridiales;f__Ruminococcaceae;g__Ethanoligenens;s__;d__denovo347 |

| | | |
|---|---|---|
| | denovo1756 | k__Bacteria;p__Firmicutes;c__Clostridia;o__Clostridiales;f__Lachnospiraceae;g__Dorea;s__;d__denovo1756 |
| | denovo3055 | k__Bacteria;p__Firmicutes;c__Clostridia;o__Clostridiales;f__Ruminococcaceae;g__Faecalibacterium;s__;d__denovo3055 |
| | denovo1779 | k__Bacteria;p__Firmicutes;c__Clostridia;o__Clostridiales;f__Lachnospiraceae;g__;s__;d__denovo1779 |
| | denovo2753 | k__Bacteria;p__Firmicutes;c__Clostridia;o__Clostridiales;f__Ruminococcaceae;g__Ethanoligenens;s__;d__denovo2753 |
| | denovo1999 | k__Bacteria;p__Firmicutes;c__Clostridia;o__Clostridiales;f__Lachnospiraceae;g__;s__;d__denovo1999 |

**Additional file 14: Table S2 a**

Correlation of species in the *Lachnospiraceae* family with CDI case that were selected by the Boruta algorithm

| Species | Correlation with CDI case |
|---|---|
| denovo413 | 0.002 |
| denovo295 | 0.006 |
| denovo545 | 0.008 |
| denovo2806 | 0.012 |
| denovo1784 | 0.016 |
| denovo913 | 0.021 |
| denovo202 | 0.021 |
| denovo3352 | 0.022 |
| denovo351 | 0.025 |
| denovo285 | 0.025 |
| denovo2947 | 0.026 |
| denovo2380 | 0.028 |
| denovo851 | 0.030 |
| denovo807 | 0.039 |
| denovo1704 | 0.039 |
| denovo2089 | 0.042 |
| denovo983 | 0.048 |
| denovo1169 | 0.050 |
| denovo1783 | 0.050 |
| denovo1048 | 0.057 |
| denovo1929 | 0.057 |
| denovo3837 | 0.062 |
| denovo2069 | 0.062 |
| denovo2960 | 0.063 |

| | |
|---|---|
| denovo458 | 0.063 |
| denovo2389 | 0.072 |
| denovo1136 | 0.073 |
| denovo2961 | 0.075 |
| denovo3923 | 0.076 |
| denovo2609 | 0.080 |
| denovo216 | 0.080 |
| denovo2297 | 0.083 |
| denovo2578 | 0.086 |
| denovo2914 | 0.089 |
| denovo2838 | 0.089 |
| denovo907 | 0.094 |
| denovo1128 | 0.095 |
| denovo2676 | 0.096 |
| denovo99 | 0.097 |
| denovo152 | 0.099 |
| denovo962 | 0.102 |
| denovo3514 | 0.104 |
| denovo1968 | 0.105 |
| denovo2720 | 0.109 |
| denovo3125 | 0.111 |
| denovo1519 | 0.112 |
| denovo3226 | 0.112 |
| denovo2289 | 0.114 |
| denovo1468 | 0.119 |
| denovo511 | 0.122 |
| denovo1865 | 0.122 |
| denovo1773 | 0.124 |
| denovo2937 | 0.132 |
| denovo104 | 0.133 |
| denovo3510 | 0.145 |
| denovo2957 | 0.145 |
| denovo150 | 0.149 |
| denovo3100 | 0.149 |
| denovo1786 | 0.151 |
| denovo3062 | 0.154 |
| denovo770 | 0.163 |
| denovo3462 | 0.166 |
| denovo2939 | 0.168 |
| denovo209 | 0.168 |
| denovo1399 | 0.186 |
| denovo3160 | 0.189 |

| | |
|---|---|
| denovo3044 | 0.190 |
| denovo1436 | 0.195 |
| denovo193 | 0.211 |
| denovo630 | 0.212 |
| denovo263 | 0.233 |
| denovo1852 | 0.235 |
| denovo595 | 0.240 |
| denovo11 | 0.298 |

**Additional file 14: Table S2 b**

Correlation of species in the *Lachnospiraceae* family with healthy controls that were selected by the Boruta algorithm

| Species | Correlation with non-diarrheal control |
|---|---|
| denovo1309 | 0.018 |
| denovo887 | 0.018 |
| denovo37 | 0.046 |
| denovo93 | 0.049 |
| denovo818 | 0.052 |
| denovo3901 | 0.053 |
| denovo931 | 0.055 |
| denovo1419 | 0.056 |
| denovo3039 | 0.064 |
| denovo3174 | 0.069 |
| denovo229 | 0.080 |
| denovo1792 | 0.094 |
| denovo932 | 0.096 |
| denovo2453 | 0.106 |
| denovo387 | 0.123 |
| denovo335 | 0.126 |
| denovo707 | 0.130 |
| denovo3762 | 0.131 |
| denovo22 | 0.137 |
| denovo377 | 0.140 |
| denovo544 | 0.151 |
| denovo2344 | 0.154 |
| denovo243 | 0.183 |
| denovo109 | 0.196 |
| denovo70 | 0.198 |

| | |
|---|---|
| denovo504 | 0.212 |
| denovo853 | 0.213 |
| denovo232 | 0.228 |
| denovo363 | 0.232 |
| denovo1295 | 0.238 |
| denovo279 | 0.242 |
| denovo479 | 0.243 |
| denovo1397 | 0.251 |
| denovo318 | 0.253 |
| denovo1359 | 0.254 |
| denovo1977 | 0.259 |
| denovo1239 | 0.261 |
| denovo724 | 0.269 |
| denovo262 | 0.271 |
| denovo1379 | 0.272 |
| denovo465 | 0.274 |
| denovo712 | 0.275 |
| denovo542 | 0.278 |
| denovo1896 | 0.280 |
| denovo1007 | 0.288 |
| denovo1228 | 0.288 |
| denovo732 | 0.290 |
| denovo903 | 0.293 |
| denovo274 | 0.294 |
| denovo940 | 0.294 |
| denovo322 | 0.301 |
| denovo371 | 0.301 |
| denovo137 | 0.306 |
| denovo445 | 0.307 |
| denovo520 | 0.308 |
| denovo411 | 0.313 |
| denovo1172 | 0.314 |
| denovo655 | 0.316 |
| denovo346 | 0.319 |
| denovo161 | 0.321 |
| denovo329 | 0.321 |
| denovo253 | 0.324 |
| denovo1173 | 0.324 |
| denovo50 | 0.328 |
| denovo258 | 0.334 |
| denovo447 | 0.336 |
| denovo414 | 0.340 |

| | |
|---|---|
| denovo289 | 0.345 |
| denovo884 | 0.349 |
| denovo301 | 0.349 |
| denovo766 | 0.352 |
| denovo74 | 0.361 |
| denovo425 | 0.366 |
| denovo84 | 0.373 |
| denovo415 | 0.374 |
| denovo422 | 0.388 |
| denovo83 | 0.388 |
| denovo275 | 0.389 |
| denovo132 | 0.392 |
| denovo348 | 0.401 |
| denovo124 | 0.408 |
| denovo90 | 0.410 |
| denovo466 | 0.421 |
| denovo111 | 0.422 |
| denovo164 | 0.423 |
| denovo349 | 0.434 |
| denovo78 | 0.437 |
| denovo196 | 0.438 |
| denovo210 | 0.441 |
| denovo113 | 0.442 |
| denovo531 | 0.442 |
| denovo165 | 0.457 |
| denovo51 | 0.458 |
| denovo272 | 0.458 |
| denovo149 | 0.464 |
| denovo129 | 0.466 |
| denovo26 | 0.476 |
| denovo121 | 0.477 |
| denovo116 | 0.478 |
| denovo324 | 0.479 |
| denovo502 | 0.486 |
| denovo378 | 0.486 |
| denovo48 | 0.495 |
| denovo284 | 0.502 |
| denovo197 | 0.507 |
| denovo182 | 0.509 |
| denovo135 | 0.525 |
| denovo49 | 0.545 |
| denovo292 | 0.563 |

| | |
|---|---|
| denovo67 | 0.569 |
| denovo61 | 0.583 |
| denovo41 | 0.589 |
| denovo56 | 0.589 |
| denovo64 | 0.590 |
| denovo103 | 0.593 |
| denovo146 | 0.603 |
| denovo44 | 0.639 |
| denovo85 | 0.669 |
| denovo43 | 0.703 |
| denovo17 | 0.713 |

**Additional file 15: Table S3**

Correlation of species in the *Ruminococcaceae* family with healthy controls that were selected by the Boruta algorithm

| Species | Correlation with non-diarrheal control |
|---|---|
| denovo1939 | 0.059 |
| denovo2273 | 0.059 |
| denovo1636 | 0.111 |
| denovo2653 | 0.123 |
| denovo1164 | 0.137 |
| denovo790 | 0.145 |
| denovo449 | 0.169 |
| denovo605 | 0.176 |
| denovo2117 | 0.177 |
| denovo1891 | 0.178 |
| denovo244 | 0.181 |
| denovo2347 | 0.185 |
| denovo320 | 0.186 |
| denovo1532 | 0.191 |
| denovo1916 | 0.204 |
| denovo1138 | 0.205 |
| denovo728 | 0.213 |
| denovo1679 | 0.214 |
| denovo593 | 0.220 |
| denovo344 | 0.223 |
| denovo1637 | 0.232 |
| denovo1269 | 0.232 |

| | |
|---|---|
| denovo347 | 0.240 |
| denovo569 | 0.245 |
| denovo740 | 0.254 |
| denovo1737 | 0.255 |
| denovo315 | 0.262 |
| denovo993 | 0.264 |
| denovo1380 | 0.264 |
| denovo543 | 0.269 |
| denovo101 | 0.269 |
| denovo356 | 0.274 |
| denovo1098 | 0.276 |
| denovo1549 | 0.276 |
| denovo510 | 0.277 |
| denovo186 | 0.295 |
| denovo1297 | 0.299 |
| denovo970 | 0.304 |
| denovo880 | 0.315 |
| denovo726 | 0.321 |
| denovo793 | 0.322 |
| denovo102 | 0.329 |
| denovo221 | 0.330 |
| denovo578 | 0.337 |
| denovo304 | 0.338 |
| denovo218 | 0.346 |
| denovo663 | 0.375 |
| denovo338 | 0.380 |
| denovo156 | 0.385 |
| denovo276 | 0.388 |
| denovo169 | 0.394 |
| denovo380 | 0.399 |
| denovo114 | 0.411 |
| denovo115 | 0.413 |
| denovo112 | 0.460 |
| denovo175 | 0.461 |
| denovo189 | 0.477 |
| denovo38 | 0.478 |
| denovo440 | 0.483 |
| denovo463 | 0.484 |
| denovo333 | 0.486 |
| denovo242 | 0.491 |
| denovo432 | 0.509 |
| denovo76 | 0.517 |

| | |
|---|---|
| denovo80 | 0.526 |
| denovo30 | 0.579 |
| denovo9 | 0.640 |
| denovo71 | 0.645 |

**Additional file 16: Table S4**

Correlation of species in the *Bacteroidaceae* family with healthy controls that were selected by the Boruta algorithm

| Species | Correlation with non-diarrheal control |
|---|---|
| denovo2100 | 0.002 |
| denovo3232 | 0.014 |
| denovo2605 | 0.015 |
| denovo2679 | 0.016 |
| denovo1775 | 0.017 |
| denovo2406 | 0.029 |
| denovo2813 | 0.029 |
| denovo3119 | 0.031 |
| denovo3614 | 0.033 |
| denovo2842 | 0.043 |
| denovo3241 | 0.049 |
| denovo1217 | 0.051 |
| denovo46 | 0.052 |
| denovo2263 | 0.056 |
| denovo2138 | 0.056 |
| denovo2382 | 0.058 |
| denovo3178 | 0.064 |
| denovo1894 | 0.065 |
| denovo1760 | 0.065 |
| denovo1123 | 0.066 |
| denovo1920 | 0.068 |
| denovo1970 | 0.069 |
| denovo2125 | 0.073 |
| denovo2437 | 0.073 |
| denovo1433 | 0.075 |
| denovo1893 | 0.080 |
| denovo1598 | 0.087 |
| denovo1180 | 0.091 |
| denovo2024 | 0.092 |

| | |
|---|---|
| denovo1044 | 0.093 |
| denovo2103 | 0.093 |
| denovo1500 | 0.093 |
| denovo1156 | 0.097 |
| denovo2008 | 0.098 |
| denovo1111 | 0.099 |
| denovo1675 | 0.101 |
| denovo1462 | 0.103 |
| denovo1903 | 0.104 |
| denovo1176 | 0.104 |
| denovo1413 | 0.105 |
| denovo1122 | 0.107 |
| denovo1298 | 0.107 |
| denovo1382 | 0.107 |
| denovo1215 | 0.110 |
| denovo2502 | 0.111 |
| denovo868 | 0.112 |
| denovo1804 | 0.115 |
| denovo1197 | 0.115 |
| denovo969 | 0.117 |
| denovo1344 | 0.118 |
| denovo1567 | 0.120 |
| denovo885 | 0.123 |
| denovo2670 | 0.124 |
| denovo644 | 0.128 |
| denovo1557 | 0.129 |
| denovo1927 | 0.130 |
| denovo1258 | 0.131 |
| denovo2223 | 0.133 |
| denovo1560 | 0.134 |
| denovo912 | 0.137 |
| denovo1662 | 0.138 |
| denovo1049 | 0.138 |
| denovo1036 | 0.139 |
| denovo1474 | 0.141 |
| denovo965 | 0.141 |
| denovo660 | 0.143 |
| denovo1994 | 0.143 |
| denovo1565 | 0.145 |
| denovo1181 | 0.147 |
| denovo1282 | 0.150 |
| denovo1192 | 0.150 |

| | |
|---|---|
| denovo1350 | 0.154 |
| denovo39 | 0.157 |
| denovo1000 | 0.160 |
| denovo1603 | 0.160 |
| denovo1271 | 0.160 |
| denovo687 | 0.161 |
| denovo838 | 0.161 |
| denovo1278 | 0.161 |
| denovo1523 | 0.164 |
| denovo1431 | 0.164 |
| denovo783 | 0.164 |
| denovo1080 | 0.166 |
| denovo973 | 0.168 |
| denovo928 | 0.169 |
| denovo1802 | 0.170 |
| denovo900 | 0.171 |
| denovo624 | 0.173 |
| denovo1423 | 0.173 |
| denovo1415 | 0.173 |
| denovo1574 | 0.174 |
| denovo950 | 0.175 |
| denovo776 | 0.175 |
| denovo870 | 0.175 |
| denovo1034 | 0.177 |
| denovo1040 | 0.178 |
| denovo565 | 0.180 |
| denovo1711 | 0.183 |
| denovo1162 | 0.183 |
| denovo47 | 0.184 |
| denovo952 | 0.184 |
| denovo844 | 0.185 |
| denovo2272 | 0.186 |
| denovo665 | 0.187 |
| denovo998 | 0.188 |
| denovo964 | 0.188 |
| denovo945 | 0.188 |
| denovo1774 | 0.189 |
| denovo774 | 0.190 |
| denovo473 | 0.190 |
| denovo536 | 0.191 |
| denovo781 | 0.192 |
| denovo1158 | 0.192 |

| | |
|---|---|
| denovo832 | 0.192 |
| denovo765 | 0.193 |
| denovo1155 | 0.194 |
| denovo2115 | 0.196 |
| denovo803 | 0.196 |
| denovo1030 | 0.198 |
| denovo936 | 0.199 |
| denovo718 | 0.201 |
| denovo611 | 0.202 |
| denovo672 | 0.203 |
| denovo1276 | 0.204 |
| denovo1692 | 0.205 |
| denovo1150 | 0.205 |
| denovo2345 | 0.205 |
| denovo682 | 0.206 |
| denovo1149 | 0.207 |
| denovo464 | 0.207 |
| denovo23 | 0.207 |
| denovo556 | 0.208 |
| denovo562 | 0.208 |
| denovo893 | 0.209 |
| denovo19 | 0.209 |
| denovo539 | 0.209 |
| denovo659 | 0.209 |
| denovo775 | 0.209 |
| denovo136 | 0.210 |
| denovo763 | 0.212 |
| denovo1113 | 0.212 |
| denovo1641 | 0.212 |
| denovo878 | 0.213 |
| denovo958 | 0.214 |
| denovo1175 | 0.214 |
| denovo1605 | 0.215 |
| denovo661 | 0.217 |
| denovo40 | 0.217 |
| denovo1322 | 0.217 |
| denovo1093 | 0.218 |
| denovo894 | 0.218 |
| denovo187 | 0.219 |
| denovo1120 | 0.220 |
| denovo1409 | 0.221 |
| denovo577 | 0.221 |

| | |
|---|---|
| denovo2065 | 0.221 |
| denovo640 | 0.223 |
| denovo654 | 0.223 |
| denovo470 | 0.224 |
| denovo513 | 0.224 |
| denovo521 | 0.225 |
| denovo906 | 0.225 |
| denovo600 | 0.226 |
| denovo622 | 0.227 |
| denovo480 | 0.227 |
| denovo1211 | 0.227 |
| denovo1012 | 0.227 |
| denovo566 | 0.227 |
| denovo951 | 0.228 |
| denovo145 | 0.228 |
| denovo1741 | 0.229 |
| denovo1429 | 0.230 |
| denovo1542 | 0.230 |
| denovo721 | 0.230 |
| denovo722 | 0.232 |
| denovo799 | 0.232 |
| denovo1246 | 0.233 |
| denovo760 | 0.233 |
| denovo1530 | 0.238 |
| denovo1245 | 0.238 |
| denovo532 | 0.239 |
| denovo754 | 0.239 |
| denovo1062 | 0.239 |
| denovo1137 | 0.239 |
| denovo768 | 0.240 |
| denovo1384 | 0.241 |
| denovo971 | 0.241 |
| denovo791 | 0.242 |
| denovo954 | 0.242 |
| denovo848 | 0.245 |
| denovo396 | 0.247 |
| denovo2162 | 0.247 |
| denovo1291 | 0.247 |
| denovo618 | 0.247 |
| denovo955 | 0.249 |
| denovo623 | 0.250 |
| denovo7 | 0.251 |

| | |
|---|---|
| denovo699 | 0.252 |
| denovo797 | 0.252 |
| denovo921 | 0.255 |
| denovo1126 | 0.257 |
| denovo856 | 0.258 |
| denovo2110 | 0.259 |
| denovo581 | 0.259 |
| denovo1108 | 0.259 |
| denovo1267 | 0.259 |
| denovo609 | 0.259 |
| denovo1196 | 0.260 |
| denovo1569 | 0.260 |
| denovo985 | 0.261 |
| denovo686 | 0.261 |
| denovo745 | 0.263 |
| denovo980 | 0.264 |
| denovo981 | 0.264 |
| denovo490 | 0.265 |
| denovo747 | 0.266 |
| denovo1072 | 0.266 |
| denovo991 | 0.268 |
| denovo1118 | 0.268 |
| denovo610 | 0.269 |
| denovo534 | 0.270 |
| denovo8 | 0.271 |
| denovo1159 | 0.273 |
| denovo563 | 0.273 |
| denovo780 | 0.274 |
| denovo685 | 0.275 |
| denovo579 | 0.276 |
| denovo1065 | 0.277 |
| denovo706 | 0.277 |
| denovo1014 | 0.280 |
| denovo608 | 0.281 |
| denovo529 | 0.283 |
| denovo552 | 0.283 |
| denovo996 | 0.284 |
| denovo670 | 0.284 |
| denovo558 | 0.288 |
| denovo1151 | 0.288 |
| denovo934 | 0.288 |
| denovo251 | 0.288 |

| | |
|---|---|
| denovo800 | 0.288 |
| denovo614 | 0.289 |
| denovo949 | 0.290 |
| denovo762 | 0.291 |
| denovo852 | 0.291 |
| denovo546 | 0.293 |
| denovo840 | 0.295 |
| denovo734 | 0.299 |
| denovo861 | 0.299 |
| denovo901 | 0.300 |
| denovo517 | 0.301 |
| denovo280 | 0.302 |
| denovo683 | 0.303 |
| denovo1249 | 0.306 |
| denovo287 | 0.309 |
| denovo882 | 0.310 |
| denovo656 | 0.311 |
| denovo223 | 0.314 |
| denovo821 | 0.314 |
| denovo833 | 0.314 |
| denovo688 | 0.314 |
| denovo974 | 0.315 |
| denovo567 | 0.316 |
| denovo867 | 0.316 |
| denovo967 | 0.322 |
| denovo634 | 0.324 |
| denovo637 | 0.325 |
| denovo475 | 0.326 |
| denovo1227 | 0.330 |
| denovo5 | 0.331 |
| denovo550 | 0.331 |
| denovo343 | 0.334 |
| denovo238 | 0.335 |
| denovo373 | 0.340 |
| denovo812 | 0.343 |
| denovo433 | 0.344 |
| denovo846 | 0.345 |
| denovo1003 | 0.354 |
| denovo496 | 0.360 |
| denovo460 | 0.365 |
| denovo715 | 0.366 |
| denovo526 | 0.383 |

| | |
|---|---|
| denovo674 | 0.385 |
| denovo629 | 0.407 |
| denovo29 | 0.411 |
| denovo27 | 0.418 |
| denovo1 | 0.422 |
| denovo172 | 0.427 |
| denovo219 | 0.429 |
| denovo557 | 0.436 |
| denovo213 | 0.449 |
| denovo139 | 0.464 |
| denovo160 | 0.474 |
| denovo184 | 0.479 |
| denovo191 | 0.489 |
| denovo201 | 0.505 |
| denovo4 | 0.523 |

**Additional file 17: Table S5**

Correlation of species in the *Rikenellaceae* family with healthy controls that were selected by the Boruta algorithm

| **Species** | **Correlation with non-diarrheal control** |
|---|---|
| denovo1667 | 0.003 |
| denovo1182 | 0.004 |
| denovo3212 | 0.005 |
| denovo1659 | 0.006 |
| denovo1357 | 0.007 |
| denovo1292 | 0.008 |
| denovo924 | 0.009 |
| denovo1870 | 0.010 |
| denovo1302 | 0.011 |
| denovo1363 | 0.012 |
| denovo841 | 0.013 |
| denovo1075 | 0.014 |
| denovo1082 | 0.015 |
| denovo1147 | 0.016 |
| denovo892 | 0.017 |
| denovo720 | 0.017 |
| denovo2011 | 0.018 |

| | |
|---|---|
| denovo914 | 0.019 |
| denovo1133 | 0.020 |
| denovo805 | 0.021 |
| denovo1364 | 0.022 |
| denovo948 | 0.023 |
| denovo680 | 0.024 |
| denovo1177 | 0.025 |
| denovo1058 | 0.026 |
| denovo1511 | 0.027 |
| denovo1027 | 0.028 |
| denovo1268 | 0.029 |
| denovo357 | 0.030 |
| denovo1507 | 0.031 |
| denovo1063 | 0.032 |
| denovo1039 | 0.033 |
| denovo62 | 0.034 |
| denovo157 | 0.035 |
| denovo920 | 0.036 |
| denovo302 | 0.037 |
| denovo446 | 0.037 |
| denovo694 | 0.038 |
| denovo117 | 0.039 |
| denovo416 | 0.040 |
| denovo86 | 0.041 |
| denovo457 | 0.042 |
| denovo6 | 0.043 |
| denovo183 | 0.044 |
| denovo144 | 0.045 |

**Additional file 18: Table S6 a**

Correlation of species in the *Erysipelotrichaceae* family with healthy controls that were selected by the Boruta algorithm

| Species | Correlation with non-diarrheal control |
|---|---|
| denovo453 | 0.040 |
| denovo383 | 0.047 |
| denovo125 | 0.053 |
| denovo107 | 0.060 |

| | |
|---|---|
| denovo231 | 0.066 |
| denovo273 | 0.073 |
| denovo392 | 0.080 |
| denovo180 | 0.086 |
| denovo240 | 0.093 |
| denovo106 | 0.099 |

**Additional file 18: Table S6 b**

Correlation of species in the *Erysipelotrichaceae* family with CDI cases that were selected by the Boruta algorithm

| Species | Correlation with CDI Case |
|---|---|
| denovo298 | 0.003 |
| denovo222 | 0.005 |
| denovo559 | 0.007 |
| denovo2197 | 0.009 |
| denovo3316 | 0.012 |
| denovo1747 | 0.014 |
| denovo1925 | 0.016 |
| denovo1785 | 0.018 |
| denovo2663 | 0.020 |
| denovo3542 | 0.022 |
| denovo2360 | 0.024 |
| denovo33 | 0.026 |
| denovo31 | 0.028 |
